# DeePEn-A Depth sensitive benchmark for Protein Engineering

**DOI:** 10.64898/2026.06.09.731073

**Authors:** Robert Schmirler, Maurice Brenner, Sebastian Franz, Michael Heinzinger, Burkhard Rost

**Affiliations:** TUM (Technical University of Munich), School of Computation, Information and Technology (CIT), Faculty of Informatics, Chair of Bioinformatics & Computational Biology - i12, Boltzmannstr. 3, 85748 Garching/Munich, Germany; AbbVie Deutschland GmbH & Co. KG, Innovation Center, BTS IR LU, Ludwigshafen, 67061, Germany; Institute of Computational Biology, Computational Health Center, Helmholtz Munich, Munich, 85764, Germany; Institute for Advanced Study (TUM-IAS), Lichtenbergstr. 2a, 85748 Garching/Munich, Germany & TUM School of Life Sciences Weihenstephan (WZW), Alte Akademie 8, Freising, Germany

## Abstract

Recent progress in modeling techniques and high-throughput screening has significantly enhanced the accessibility of protein engineering. Nevertheless, further progress gets hindered by the lack of robust benchmarks that capture the practical challenges for real-world protein engineering. Here, we introduced DeePEn, a **De**pth-s**e**nsitive benchmark for **P**rotein **En**gineering that quantifies model generalization to held-out variants at increasing mutational distance from the wild type. We defined distance as the number of simultaneous point mutations, i.e., single amino acid variants (SAVs), moving from wild-type to mutant (*edit distance* in computer science jargon). DeePEn retrospectively evaluates prediction and prioritization using experimentally labelled variants from deep mutational scanning (DMS) datasets; it does not benchmark prospective sequence generation or goal-directed optimization. Specifically selecting four DMS datasets with sufficient multi- mutation data points from ProteinGym, we assessed recent predictive models, including general and biophysics-informed protein Language Models (pLMs), and a non-transformer neural network. Our results highlight how the performance of all models deteriorates with increasing mutational distance and that no single metric sufficiently captures the diverse requirements of protein engineering. To overcome these shortcomings, DeePEn provides a readily available resource for multi-metric benchmarking that focuses on variant-effect prediction of variants distant from the wild type.

## 1. Introduction

Deep mutational scanning (DMS) comprehensively gauges the impact of sequence changes [1–4] often referred to as the protein function fitness landscape. The excellent *ProteinGym* [1] resource collects and curates DMS datasets (217 in all 05/2026), for assessing performance of methods predicting the impact of sequence variants. However, the majority of ProteinGym’s DMS datasets (148 of 217) contain exclusively single amino acid variants (SAVs), i.e., the change of one single amino acid at a time, also dubbed point mutation. For a protein with L residues, L*19 is the maximal number of such SAVs. The remaining 69 datasets (Fig. 1a) contain both SAVs and higher order variants (with more than one mutation) where the number of possibilities rises exp onentially, with N_d_ as the number of possible variants at distance d, i.e., the number of consecutive SAVs (also referred to as the mutational depth; (Eq. 1)[5].

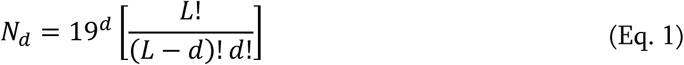

**Fig. 1:**
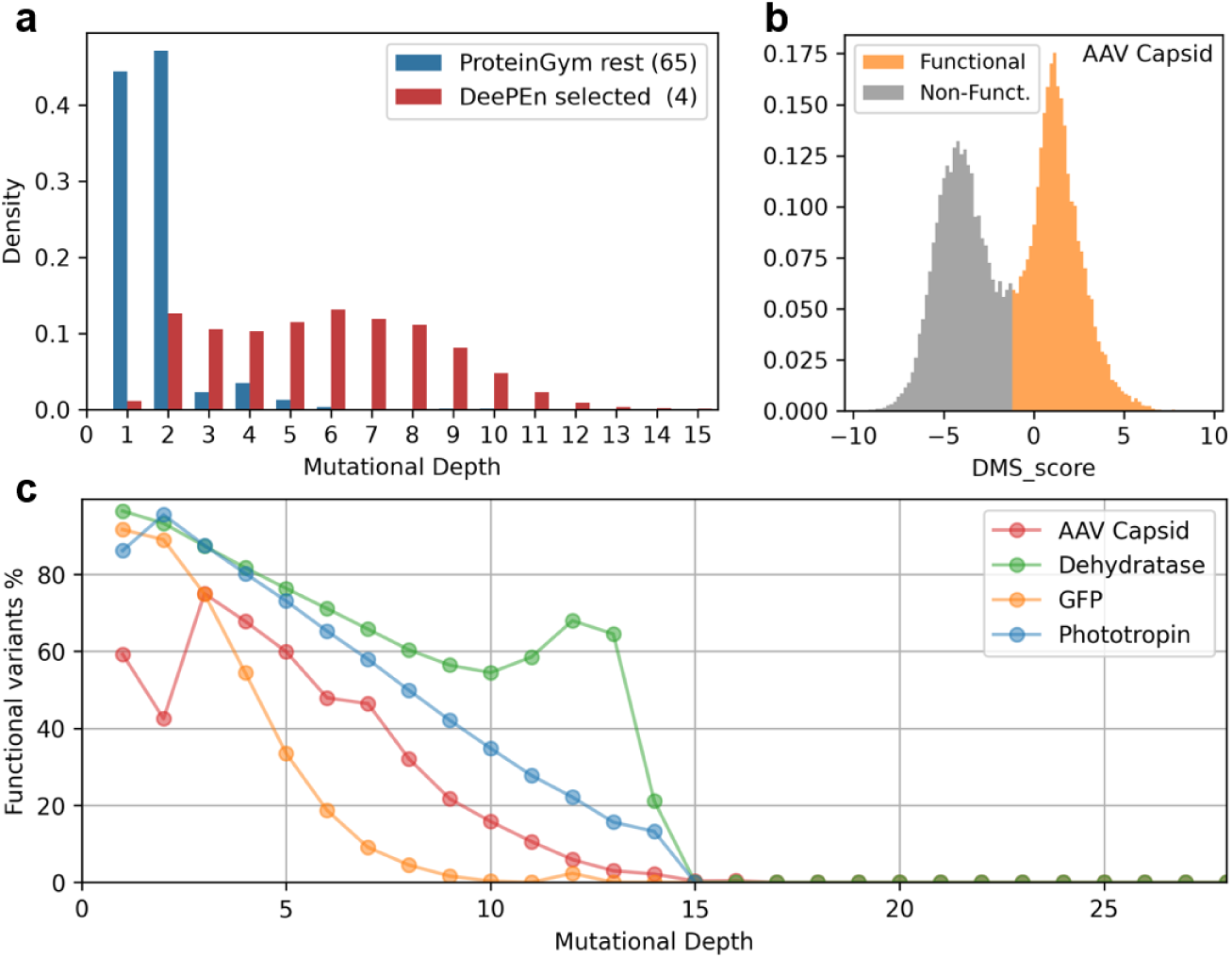
Experimental data in ProteinGym and its DeePEn subset. Panel. **a** shows the average distribution of sequence variants for the 69 datasets in *ProteinGym* [1] which contain higher order variants. Mutational depth on the x-axis is defined as the number of simultaneous single Amino Acid Variants (SAVs). For instance, depth=6 contains sequences in which six residues differ from wild-type. The colors differentiate between *Selected*, i.e., the four data sets covered by our DeePEn collection, and *Other* as all the other 65 (69-4) sets. Most of the latter datasets were described by a power law distribution with fitness data mostly available for single and double mutants. There was only a negligible number of variants beyond depth 15. **Panel b** provides the distribution of fitness scores (x-axis: DMS_score representing the experimental value) as well as their binary classification (taken from the original publication [6]) for the *AVV Capsid* dataset. **Panel c**: In all four selected DMS datasets the fraction of functional variants (y-axis: fraction of functional variants as percentage) decreases with increasing depth (x-axis: mutational depth as described in panel a). Note that this decrease is trivial for experiments randomly changing all possible SAVs at all depth, e.g., if at d=1 there were 50% functional SAVs, a depth d=10 this would have shrunk to 1/(2^10) roughly 0.1%. Due to the clever design of DMS experiments, the observed values are substantially higher (by almost 4 orders for the Dehydratase).

In contrast to the space of exploding possibilities, the vast number of variants annotated by DMS experiments are of distance d≤2 (Fig. 1a). We therefore selected the small subset from ProteinGym with sufficient depth and data set size (details in SOM, Supporting Online Material, Table S1) which left us with four sets (of 217, Methods, Table 2) for our **De**pth s**e**nsitive **P**rotein **En**gineering benchmark (DeePEn). Throughout this work, we refer to each of those by the common name of the protein used in their DMS experiments:

- “*AAV Capsid*”, measuring capsid assembly and genome-packaging fitness of AAV2 VP1 in a diverse dataset using multiple sampling strategies spanning single substitutions, randomly selected multi-mutants, and ML-guided designs [6].
- “Dehydratase”, measuring growth rate conferred by yeast His3 (an enzyme essential for histidine synthesis), in a dataset generated using 12 independent gene-segment libraries that recombine amino acid states found in extant yeast orthologues [7].
- “GFP”, measuring green fluorescence intensity of Aequorea victoria GFP, in a dataset generated by random mutagenesis of the GFP coding sequence [8]
- “Phototropin”, measuring cyan-green fluorescence intensity of the oxygen-independent FMN-binding fluorescent protein CreiLOV, derived from the LOV domain of *Chlamydomonas reinhardtii* phototropin, in a combinatorial dataset generated by combining 20 mutations at 15 sites selected through single-site saturation mutagenesis [9].

**Table 1:** DeePEn results - normalized metric values ^Δ^.

| *Model* | *Depth* | *AUCPR<sub>norm</sub>* | *Spearman<sub>func</sub>* | *TOP10%<sub>func</sub><sub>norm</sub>* | *NDCG@10%<sub>func</sub><sub>norm</sub>* |
|---|---|---|---|---|---|
| *Perfect classifier baseline* | *shallow (5-9)* | 100 ± 0 | 0 ± 0 | 5 ± 0 | 62 ± 0 |
|  | *higher depth (10+)* | 100 ± 0 | 0 ± 0 | 7 ± 0 | 69 ± 0 |
| *SBI baseline* | *shallow (5-9)* | 37 ± 0 | 21 ± 0 | 10 ± 0 | 36 ± 0 |
|  | *higher depth (10+)* | 15 ± 0 | 12 ± 0 | 3 ± 0 | 15 ± 0 |
| *1-hot baseline* | *shallow (5-9)* | 15 ± 19 | 2 ± 12 | 1 ± 5 | 7 ± 30 |
|  | *higher depth (10+)* | 11 ± 14 | 4 ± 19 | 1 ± 5 | -3 ± 28 |
| *ESM2_650M<sub>zero-shot</sub>* | *shallow (5-9)* | 39 ± 0 | 22 ± 0 | 13 ± 0 | 48 ± 0 |
|  | *higher depth (10+)* | 52 ± 0 | 34 ± 0 | 17 ± 0 | 57 ± 0 |
| *Tranception_L<sub>zero-shot</sub>* | *shallow (5-9)* | 56 ± 0 | 27 ± 0 | 13 ± 0 | 64 ± 0 |
|  | *higher depth (10+)* | 48 ± 0 | 30 ± 0 | 15 ± 0 | 63 ± 0 |
| *MAVE-NN_add* | *shallow (5-9)* | 75 ± 1 | 42 ± 2 | 26 ± 2 | 77 ± 1 |
|  | *higher depth (10+)* | 30 ± 3 | 25 ± 1 | 22 ± 2 | 48 ± 3 |
| *MAVE-NN_bb* | *shallow (5-9)* | 75 ± 11 | 45 ± 18 | 27 ± 12 | 70 ± 14 |
|  | *higher depth (10+)* | 42 ± 13 | 28 ± 19 | 20 ± 8 | 40 ± 12 |
| *METL-global_1D* | *shallow (5-9)* | 76 ± 2 | 47 ± 3 | 28 ± 4 | 74 ± 6 |
|  | *higher depth (10+)* | 51 ± 4 | 31 ± 7 | 19 ± 6 | 47 ± 10 |
| *METL-global_3D* | *shallow (5-9)* | 73 ± 3 | 48 ± 3 | 28 ± 3 | 72 ± 6 |
|  | *higher depth (10+)* | 57 ± 7 | 35 ± 6 | 25 ± 6 | 60 ± 9 |
| *PT5-LoRA_reg* | *shallow (5-9)* | 81 ± 5 | 53 ± 4 | **34 ± 7** | 76 ± 11 |
|  | *higher depth (10+)* | 65 ± 16 | 36 ± 8 | 25 ± 13 | 57 ± 27 |
| *PT5-LoRA_rank* | *shallow (5-9)* | 82 ± 3 | **54 ± 4** | 33 ± 7 | **82 ± 4** |
|  | *higher depth (10+)* | **68 ± 15** | **39 ± 13** | **27 ± 10** | **69 ± 25** |
| *PT5-Emb* | *shallow (5-9)* | 78 ± 4 | 46 ± 5 | 25 ± 5 | 74 ± 6 |
|  | *higher depth (10+)* | 61 ± 10 | 35 ± 9 | 19 ± 12 | 56 ± 25 |
| *PT5-Emb_evo* | *shallow (5-9)* | **83 ± 4** | 49 ± 4 | 28 ± 7 | 78 ± 8 |
|  | *higher depth (10+)* | 67 ± 12 | 32 ± 20 | 21 ± 16 | 65 ± 28 |

<sup>Δ</sup> Results per model for both depth bins (shallow/higher) give averages over all depth subsets and all four datasets. Baseline methods on top, supervised methods below. To control for shifting class prevalence, metrics are normalized (per depth subset, Eq. 5) to a range between 0 (random) and 1 (perfect prediction). Spearman<sub>func</sub> is not affected as its random performance is already 0. Absolute values can be found in SOM Table S2. 95% confidence intervals of the mean values were only calculated for model replicates within individual depths/dataset-combinations. Metric values multiplied by 100, numerically best performing model per depth and metric in bold, these model differences are not statistically significant. The Perfect classifier baseline is perfectly separating non-functional and functional variants while randomly shuffling all functional variants.

**Table 2:** DeePEn datasets.

| Dataset | Split | Distance from wildtype | Number of seq | Number of functional seq | Share of functional seq |
|---|---|---|---|---|---|
| AAV capsid | Train | 1-3 | 18,271 | 10,093 | 55.2% |
|  | Valid | 4 | 6,646 | 4,504 | 67.8% |
|  | Test | 5-9 | 13,942 | 7,132 | 51.2% |
|  |  | 10-28 | 3,469 | 183 | 5.3% |
| Dehydratase | Train | 1-3 | 9,270 | 8,198 | 88.4% |
|  | Valid | 4 | 25,927 | 21,177 | 81.7% |
|  | Test | 5-9 | 429,661 | 283,521 | 66.0% |
|  |  | 10-28 | 31,279 | 17,308 | 55.3% |
| GFP | Train | 1-3 | 26,197 | 21,594 | 82.4% |
|  | Valid | 4 | 9,387 | 5,104 | 54.4% |
|  | Test | 5-7 | 13,649 | 3,315 | 24.3% |
|  |  | 8-15<sup>*</sup> | 2,481 | 73 | 2.9% |
| Phototropin | Train | 1-3 | 3,276 | 2,852 | 87.1% |
|  | Valid | 4 | 3,565 | 2,859 | 80.2% |
|  | Test | 5-9 | 123,206 | 66,552 | 54.0% |
|  |  | 10-15 | 37,482 | 11,505 | 30.7% |

<sup>\*</sup>.to retain sufficient statistical power and avoid excluding GFP from the “higher depth” category of the benchmark (due to the lack of functional sequences), we decided to extend the deepest evaluation bin to a range of 8-15 for this dataset. Details can be found in SOM Table S4.

This study introduces a concise benchmark for variants of increasing distance from the wildtype, i.e., of increasing mutational depth. Considering variants at increasing distance from wildtype is important because reaching multiple, potentially conflicting, design goals often require the accumulation of multiple compatible beneficial mutations. Residue positions that could affect molecular protein function - and thereby also impact the biological process – are most relevant for typical in silico protein engineering. Thus, we focus on identifying and ranking such functional protein variants.

Our benchmark evaluates variant-effect predictors that score and prioritize candidate sequences. Such predictors can serve as components - in this context often dubbed oracles - within protein-engineering workflows, including reinforcement learning [10,11], Bayesian optimization [12,13], flow matching [14,15]. DeePEn only evaluates this predictive component retrospectively on held-out DMS variants; it does not evaluate the generation strategy or the outcome of a prospective optimization campaign.

Although our subset DeePEn clearly over-represents deep mutants compared to all sets arduously collected by ProteinGym, even in our tiny subset immensely under-represents the true spectrum (Eqn. 1; Fig. 1a: blue vs. red). All variants in *ProteinGym* come with a binary label that splits the data into functional and non-functional variants (also referred as fit/unfit). When available, ProteinGym took a biologically motivated threshold for binarizing directly from the original publications. If not (for the DeePEn subset this applied to Phototropin), ProteinGym used the median value as threshold. The data classified as functional tends to be more valuable for protein engineering (Fig. 1b as an example). Another typical pattern in functional landscape data is the strong decrease in functional variants with increasing mutation depth (Fig. 1c: toward the right). This implies that datasets are biased towards non-functional variants at higher depth. This is crucial to select suitable metrics for model evaluation.

## 2. Results

### Fine-tuning foundation pLMs captured fitness

The *Phototropin* dataset (Fig. 2) has been shown to be of high quality and assay reproducibility [9]. Therefore, we used it to determine a suitable depth cutoff for our training-, validation-, test-split. We predicted Phototropin activity inputting the foundation pLM ProtT5 (PT5) [16] with parameter-efficiently fine-tuning (PEFT) [17].

**Fig. 2:**
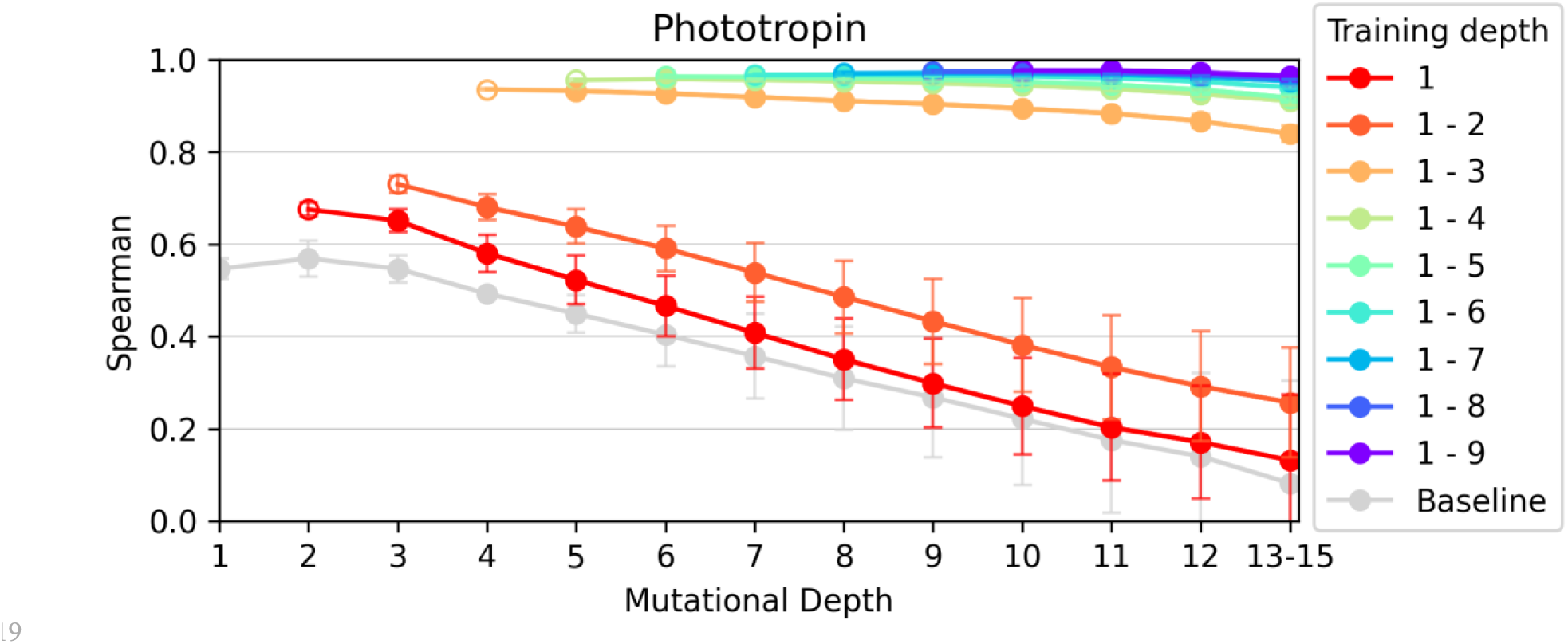
Generalizing to higher-order mutants requires training on such. Lines are colored by the depth used for training (1-n implied training contained mutations with d<(n+1)). Each point represents the Spearman correlation between predicted and actual values for a specific mutational depth. The leftmost point (for training depth 1-3 the validation is d=4) of each line was used for validation (empty circle), while all other points were the test set (full circles). Different training depth significantly impacted model capability. For the *Phototropin* dataset shown here (Fig. S1 for the other datasets), SAVs and double mutations were not sufficient to achieve high predictive performance (both red lines with training depth 1 and 1-2). When adding triplets to the training data (orange line, training depth 1-3) values increased substantially across all test depths. The Baseline was trained on 184 non-homologues non-used DMS datasets from ProteinGym (Methods).

To gauge the effect of mutation depth on model performance, we classified validation- and test- set variants by mutational depth (number of simultaneous mutations from wildtype). In all data splits, we exclusively trained on low order variants (d_val|test_>d_train_; (Fig. 2: Training Depth). We then selected all variants with one additional mutation (d_val_=d_train_+1) for validation (Fig. 2: empty circles), and the test set (Fig. 2: full circles) contained any remaining higher order depths (all with d>(d_train_+1)). In contrast to previous work applying distance-based splitting [18–20], we measured the Spearman correlation for each depth separately. Although reducing statistical significance, this avoided an undesirable bias towards the more frequent shallow (small d) variants during evaluation. Importantly, this evaluation does not reward using distance as a proxy for fitness, i.e, predicting any deep mutation as non-functional (Fig. 1c). For the highest-order variants, we always binned multiple depths to account for low counts and allowing comparisons of statistical significance (Methods, Table 2 and SOM, Tables S2-S5).

As a baseline we trained the same model on 184 out of the 213 remaining DMS sets in ProteinGym (to avoid data leakage we excluded all homologs to the DeePEn proteins, Methods). Note that this baseline is dominated by data of depth d=1, i.e., proxies the situation of the red line in Fig. 2 (Training depth: 1) apart from training on non-homologues data. While the baseline was comparable to training on single mutations of the *Phototropin* landscape, it fell short for the *GFP* and *AAV Capsid* datasets (SOM Fig. S1).

We proceeded alike for the other three datasets (SOM, Fig. S1) and chose d_train_=3 for all further experiments. Although d_train_=2 looked largely sufficient to capture interaction information in the other three datasets (SOM, Fig. S1), the low abundance of double mutants in Phototropin (176 variants; SOM Table S6) necessitated including triplets. This threshold gave a reasonable tradeoff between providing enough data to learn interactions from duplets and triplets while retaining sufficient higher-order data points (d≥5) to measure the ability of models to generalize to higher depths.

To compare supervised methods with zero-shot baselines we included ESM2_650M [21] and Tranception_L [22] results (predictions from ProteinGym). In order to explore the usefulness of data from synthetic simulation and protein structure enhanced prediction, we also compared the biophysics-informed METL [23] global models (Fig. 3, Table 1). As a non-transformer model, we investigated MAVE-NN [24].

**Fig. 3:**
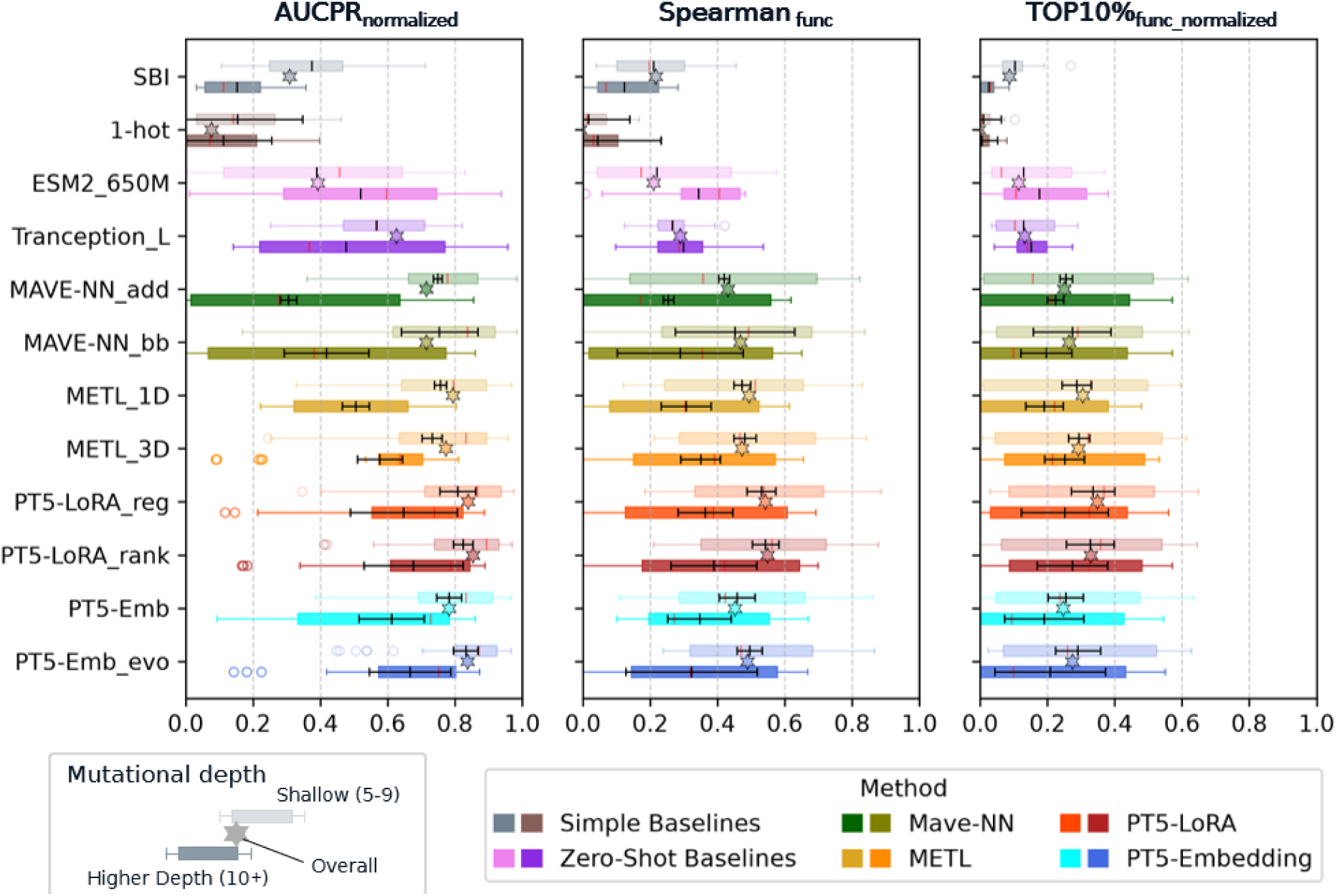
DeePEn method pooled averages, showed performance decreased with increasing depth across metrics. Two boxplots per model show the distribution of the respective metric across depth subsets (within the shallow/higher bins) across all four datasets. The star provides the average metric when calculated over the entire test set without any depth binning or subsets. Methods: in gray simple baselines: SBI - Similarity-based inference and 1-hot - as one-hot encoding (training on amino acid sequence in one-hot encoding). Zero-shot baselines in pink: ESM2_650M (additive masked log-odds) and Tranception_L (autoregressive joint likelihood). *MAVE-NN_add* uses a fixed additive effect model while the black box variant trains a neural network for effect aggregation. *PT5-LoRA* models are LoRA finetuned ProtT5 models, while *PT5-Emb* utilizes frozen ProtT5 embeddings. *METL-3D* is structure informed while *1D* is not. 95% confidence intervals of the mean values were only calculated for model replicates within individual depths/dataset-combinations, which allowed excluding any differences originating from data differences and better capturing uncertainties originating from the models. To control for shifting class prevalence, metrics are normalized (per subset, Eq. 5) to a range between 0 (random) and 1 (perfect prediction). Across models and metrics prediction performance decreased with increasing depth.

### Single metric insufficient for evaluation

Most evaluations of DMS predictions focus on the Spearman ranking correlation over the entire test set [3,18,19]. Instead, we found it necessary to compare at least four different metrics to gauge how well a method meets different ends of protein engineering. The first, AUCPR, is the standard area under the precision-recall curve as a balanced metric to gauge model ability to distinguish functional from non-functional variants (utilizing binarized scores from ProteinGym). The second, Spearman_func_ limits the computation of the Spearman correlations to the subset of functional variants instead of using the entire data. This focus reduces obfuscation from non-functional variants, which are less interesting for protein engineering. Lastly, we introduced two metrics focusing on the model’s ability to identify highly active variants. We followed previous suggestions [1] using normalized discounted cumulative gain (NDCG) [25,26], a ranking metric primarily used in information retrieval. We also introduced the simpler series of TOPn%_func_-Recall metrics for which we calculated the overlap between the subset of top n% functional variants as ranked by DMS experiments and predictions. While the addition of a free parameter *n* complicates the analysis, we considered this complication necessary because the value will depend on particular experimental objectives, i.e., will be user/task-specific. In the main body we provided results for n=10% (values for n=5, 15, 20, 25, 30 in SOM Table S13).

In analogy to our initial exploration (Fig. 2), we first evaluated each test dataset at each mutational depth separately. We then reported the average across all four datasets and subsets belonging to the “Shallow” or “Higher Depth” bins (Fig. 3, Table 1). To make sure we aggregate comparable values over depth subsets with varying class prevalence, we normalized metrics (Eq. 5) to a range between 0 (random) and 1 (perfect prediction).

The CI95 confidence intervals (points at which distributions overlap by fewer than ±1.96 standard errors) represent only model variation between replicate training runs (each model was trained three times on each dataset), not the differences between datasets and individual depth subsets.

### Models struggle to capture higher-order variant effect

Across the DeePEn datasets and all metrics, the supervised methods performed numerically worse for variants of higher depth (d≥10) than for shallow variants of shallow (5≤d≤9; Fig. 3, Table 1). The zero-shot baselines were outliers here with Tranception_L showing limited depth difference and ESM2_650M performing numerically better at higher depth. Supervised models (MAVE-NN, PT5 and METL) generally achieved numerically higher performance than the two simple baselines, similarity-based inference (SBI) and one-hot encoded sequences (1-hot). The differences between supervised models were smaller and statistically insignificant. Numerically, the PT5-LoRA_rank model (ProtT5 embeddings fine-tuned by LoRA on training set) reached the highest values (Table 1). The *MAVE-NN_add* model using additive effect modeling had the smallest variability between individual training runs (error bars Fig. 3), followed by both METL models. Our ProtT5-derived methods (PT5-*) along with the MAVE-NN and the blackbox version *MAVE-NN_bb* varied more strongly between training runs, especially for higher depth.

### Dataset differences dominated

When aggregating results from all supervised methods on a per dataset basis (Fig. 4), dataset influence became visible. Overall, the *Dehydratase* (lowest Spearman_func_ and TOP10%_func_) and *AAV Capsid* (lowest AUCPR and NDCG10%_func_) were most difficult to predict across all metrics. With *Phototropin* set being the easiest and the *GFP* set in between. Higher depth showing lower performance held here again, the only exception was TOP10%_func_ for *GFP*. The errors shown in Fig. 4 are 95% confidence intervals calculated per subset (unique depth) across models and random seeds. These narrow CI suggested that models mostly agree (extrapolate in similar fashion from the training data) and the wide box plot distribution was caused by subset/depth differences within the two depth bins. Individual dataset results per model can be found in SOM: Tables S7-S10 and Fig. S2-S5.

**Fig. 4:**
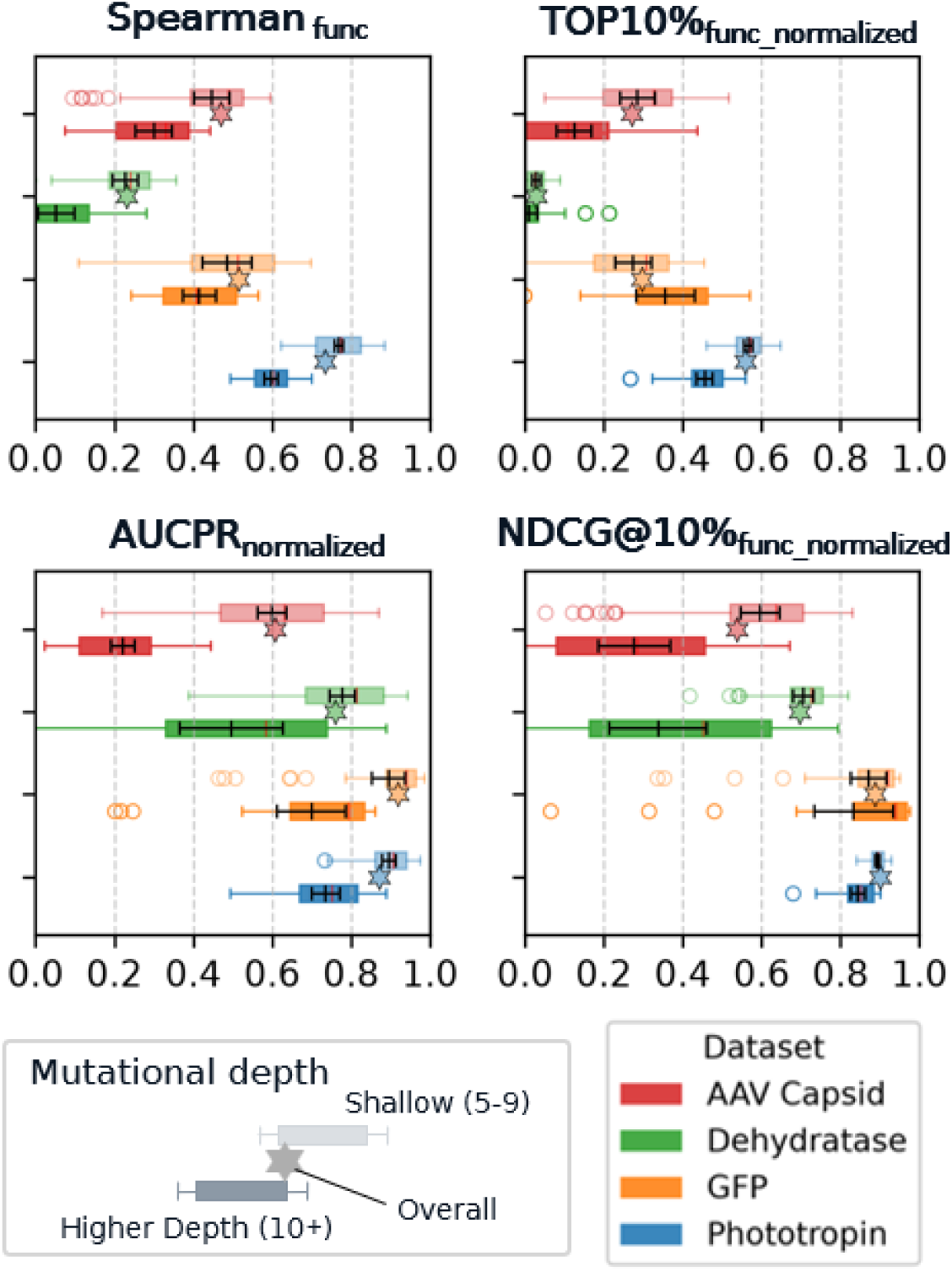
DeePEn dataset pooled supervised averages, confirmed performance decreases at higher depth. Two boxplots per dataset for different test depths show the distribution of the respective metric across depth subsets across all eight supervised methods. The star provides the average metric when calculated over the entire test set without any depth binning. 95% confidence intervals of the mean values were calculated across models and replicates within individual depths bins, which allowed excluding any differences originating from depth differences within a subset. To control for shifting class prevalence, metrics are normalized (per bin) to a range between 0 (random) and 1 (perfect prediction). Across datasets and metrics prediction performance mostly decreased with increasing depth (TOP10%_func_ for *GFP* was the exception).

### Overall test set performance skewed towards shallow bin

When comparing our depth binned results (boxplots) in both Fig. 3,4 with the same metrics simply calculated across the entire test set (stars in Fig. 3,4), these overall results were closely tracking the shallow bin performance. This was caused by the overrepresentation of shallow variants in test sets.

### Improved initialization not beneficial for fine-tuning

We applied evo-tuning [27] (unsupervised pre-training on homologs of to the protein of interest, Methods) and EVA initialization [28] to LoRa [29] weights. Neither strategy improved performance (SOM Table S11). Possibly, the major fine-tuning altered the model so much change that minor modifications to model initialization remained insignificant (SOM Fig. S6). Evo-tuning numerically improved embedding-based predictions (Fig. 3, *PT5-Emb_evo* > *PT5-Emb*), but confirming previous results, fine-tuning outperformed pretrained embeddings [17,30].

### Ranking losses superior for regression tasks

We confirmed previous findings [31] that using a ranking loss (Fig. 3: *PT5-LoRA_rank*) can improve over MSE loss (Fig. 3: *PT5-LoRA_reg*) for regression tasks (Fig. 3, Table 1). We compared different ranking losses (SOM Table S12) and selected the one with highest numerical performance: a lambda ranking loss [32] weighed by *λ* = Δ*NDCG* (Normalized Discounted Cumulative Gain, Methods [25,26]). For our PEFT [17,33,34] ranking model (Fig. 3: *PT5-LoRA_rank*) we also switched to Spearman ranking correlation for early stopping.

### Additional evaluation metrics capture diverse aspects

We created a toy dataset to help clarify the implications of different metrics. Our synthetic dataset consisted of one million random data points, of which the top 20% were labelled as *functional*. Next, we determined the true ranking of those sequences (*Perfect ranking* Fig. 5). By construction (‘perfect’), all functional variants (red) were ranked above the non-functional (blue). The *Random* baseline shuffled all ranks and for the *Balanced method* we sorted predictions such that all five metrics reached 0.7. In contrast to this balanced prediction, we designed different variations (*Methods* #1-3) for which the overall Spearman correlation and some of the remaining four metrics (Spearman_func_, AUCPR or NDCG@10%_func_ / TOP10%_func_) remained constant at 0.7, while the other metrics were substantially worse. *Method #1* demonstrated that the Spearman correlation alone can be blind to suboptimal binary classification, with an entire cluster of misplaced non-functional sequences ranked higher than the functional ones. *Method #2* highlighted the blind spot of binary classification metrics such as AUCPR: the inverted ranking within the functional sequences (the entire red color gradient is upside down) did not affect those metrics but had a dramatic effect on Spearman_func_, NDCG@10%_func_ and TOP10%_func_. Lastly *Method #3* focused on the top-heavy metrics NDCG@10%_func_ and TOP10%_func_. Those were only evaluating the very top of the ranking and therefore missed the substantially worse predictions below.

**Fig. 5:**
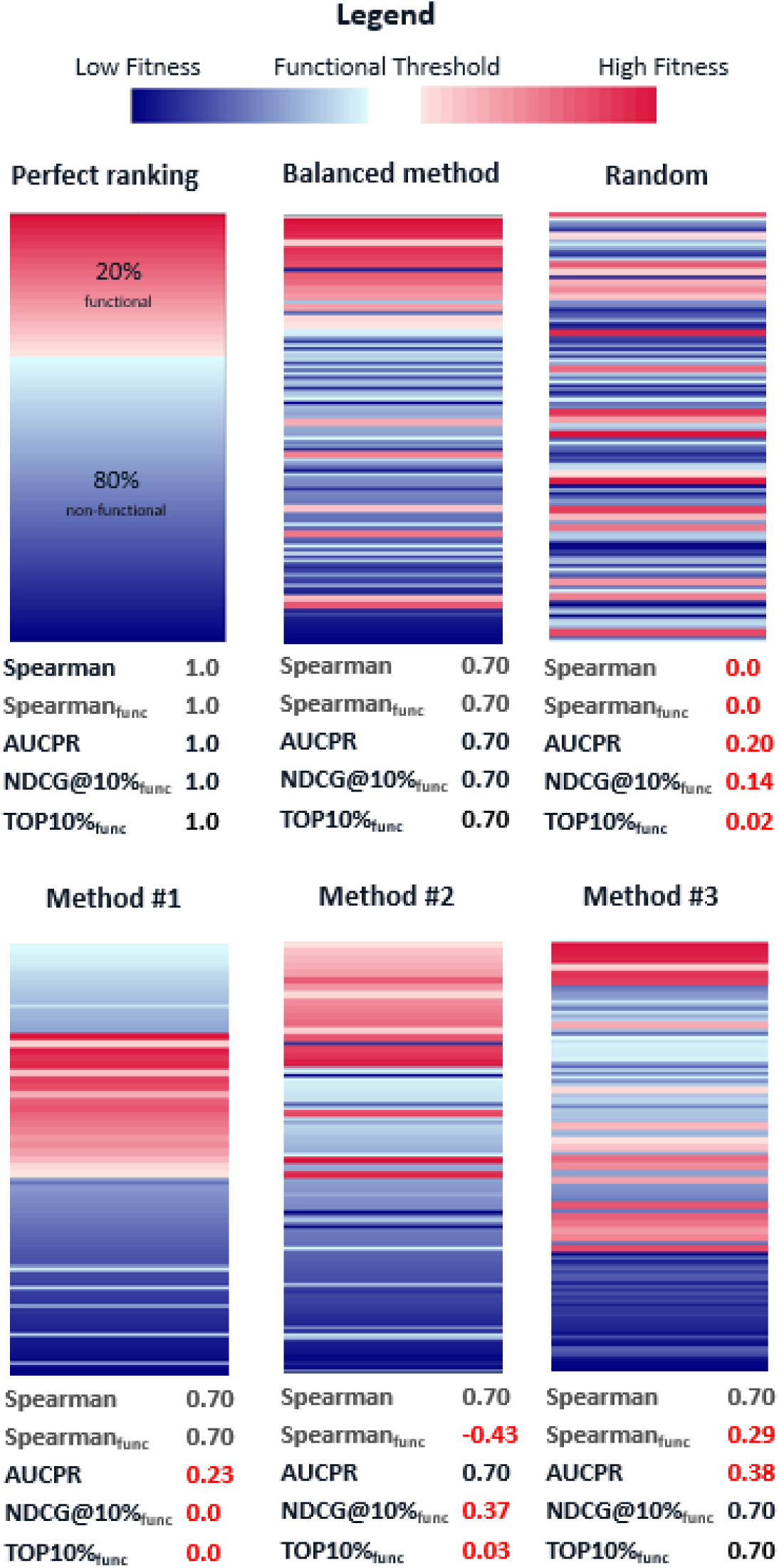
Evaluation metrics capture different aspects. A synthetic dataset with a 20:80 functional:non-functional split. Functional (red) slices are plotted with twice the height of non-functional (blue) ones for better visibility. We reported the same five evaluation metrics for each of the six hypothetical methods. The upper row shows baselines. We constructed the *Balanced method* to achieve 0.7 in all five metrics. *Random* was uniformly shuffled. We tweaked Methods #1-3 to exemplify the potential independence of different metrics: Spearman stayed constant at 0.7 throughout, while some other metrics reduced substantially. The numbers in red font indicated a decrease compared to the *Balanced method*. Reporting only Spearman would completely miss how much less desirable Methods #1-3 are compared to the *Balanced method*.

Our synthetic dataset demonstrated that the metrics capture distinct aspects of prediction quality, although the deliberately constructed examples represent a worst-case scenario. To assess their relationship in practice, we compared the model rankings produced by each metric on DeePEn (SOM Fig. S7). For both shallow and higher depth bins, model rankings were moderately to strongly correlated (Spearman between 0.6 and 0.95), indicating that models performing better on one metric often also performed better on the others.

### Low N sampling successful at shallow depth

Lastly, we investigated the potential of sampling a very small amount (a single 96 well plate) of top variants (Fig. 6) using the models tested here. This showed that models can succeed even with a small sampling size, even though it is not guaranteed (no supervised model ranked even a single top100 *Dehydratase* variant in its predicted top96).

**Fig. 6:**
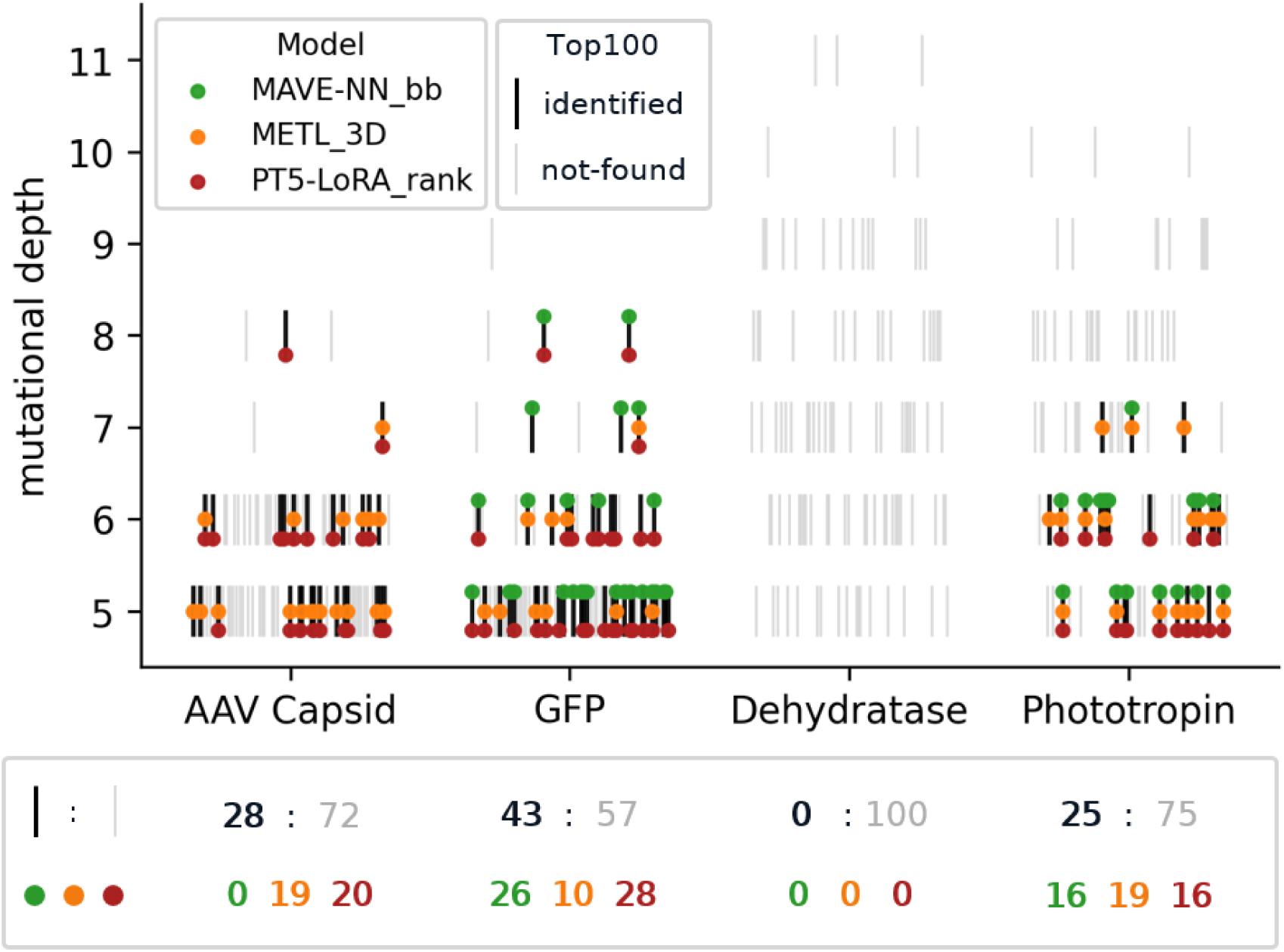
Top100 identified by 96-well plate sampling. Grey (not found) and black (identified, by at least one model) dashes each stand for one of the top100 variants per dataset. Lines are ordered by their DMS score (high to low; right to left) and assigned to their respective mutational depth (y-axis). Colored circles show the overlap with the predicted top96 (using the average rank from three training runs with different random seeds) for three models. The box below provides the number of “identified” : “not found” top100 variants, per dataset as well as the number of top100 “hits” on the simulated 96-well plates per model and dataset.

This seemed contradictory to our overall finding of models showing low TOPn%_func_ values especially for higher depth. But in fact, when looking at the overall datasets (instead of the binned version for the benchmark) most top100 variants (Fig. 6 light dashes) were shallow (especially for the *AAV Capsid* and *GFP* datasets) and successfully identified top100 variants (Fig. 6 black dashes) were even shallower (mostly depth of 5-6). For the *Phototropin* dataset, all models identified a few of the top100 variants, but only three of those were of depth 7 and none of them were further away. This clearly underrepresented the underlying distribution with 48/100 of actual top100 *Phototropin* variants having a distance greater 6.

## 3. Discussion

As a simplified illustration, assuming independent prediction errors, a method incorrectly predicting 30% of single mutations would be expected to incorrectly predict 51% (1-0.7^2^) of double mutants. This calculation is not intended as a quantitative expectation for real protein landscapes, where epistasis and correlated errors violate the independence assumption. Nevertheless, it illustrates why performance may decrease with increasing mutational depth, as observed for all supervised models across all four datasets.

A key difference from previous work [18–20] is that DeePEn evaluates each mutational depth separately rather than pooling all test variants into a single performance estimate. Without this depth subset evaluation, the depth dependent decline in predictive performance would remain hidden. Moreover, because all four test sets are dominated by more abundant shallow variants, one overestimates model performance for higher depth/deep variants, as illustrated by the overall test set values (stars) in Fig. 3 and Fig. 4. Evaluating each depth in isolation is therefore essential for measuring how reliably models extrapolate beyond shallow mutational neighborhoods.

Results aggregated over different levels of mutation depths suggested substantial performance variation (Fig. 3 boxplots). Closer inspection revealed these to originate from multiple superposing effects. Inter dataset differences (Fig. 4, SOM Fig. S2-S5, Table S7-S10) were consistent across models and dominated the other effects. The mutationally sparse *Dehydratase* dataset which was created by recombining diverse His3 orthologs [7] was the most challenging to predict. On the other end the recombination engineered *Phototropin* landscape [9] showed the best predictive performance. Intra dataset differences, i.e., depth related differences within the shallow and higher depth bins (boxplots in Fig. 4 and SOM Fig. S2-S5) also contributed. Finally, model performance varied across training runs using different random seeds. When it comes to method comparison, we consider this internal model variation the most relevant. Therefore the 95% confidence intervals reported in Fig. 3 and Table 1 were computed exclusively from random-seed variability.

While all models tended to perform worse at higher than at lower depth, some notable differences between models stood out. By design, the additive model [24] (*MAVE-NN_add*) had narrow confidence intervals (low variance between training runs with different random seeds). Other supervised models outperformed *MAVE-NN_add* at higher depths. This underlined the importance of non-additive epistatic interactions (SOM, Fig. S9). Surprisingly, the biophysically informed METL-global [23] did not improve over general pLMs when generalizing to higher depth, with little difference between the structure-informed 3D and the 1D model. As suggested by the authors, 148 proteins might not suffice for the global models to generalize. Potentially, individual METL-local models, pretrained on the four DeePEn proteins, might perform better, however, without licenses we could not investigate this. On average, zero-shot baselines did not match the performance of supervised methods, with the *Dehydratase* dataset being the exception (SOM Table S8, Fig. S3). Supervised methods faltered here due to the dataset’s mutational sparsity, where higher-order mutants lack the constituent single-mutation training data required to learn fitness effects. Conversely, zero-shot models, which inherently capture coevolution and conservation patterns, excelled because this library was generated by recombining natural segments across orthologous His3 sequences, so the models could make use of their learned evolutionary patterns.”

Fine-tuning pLMs with ranking losses numerically improved performance across all metrics (SOM Table S12) compared with standard regression training. Notably, the increase in NDCG (when using lambda ranking loss) indicated a desirable shift in predictive focus toward highly functional variants.

Variant effect prediction (VEP) evaluation might mislead when exclusively based on a single metric. Especially the overall Spearman correlation can mislead (Fig. 5). We encountered an example for this in this work. The *GFP* dataset showed a very low Spearman correlation at higher depths (7 and 8-15) in Fig. S1 (SOM), but when looking closer (SOM Table S9) the four evaluation metrics used for DeePEn did not reflect this sharp drop in performance. In this case the low overall Spearman correlation only resulted from the model’s inability to correctly rank non-functional sequences, which is largely irrelevant for engineering purposes. As non-functional variants were increasingly dominant at higher depth (SOM Table S5) this led to a sharp drop in overall Spearman for those depth.

Our constructed example (Fig. 5: Methods 1–3 vs. Balanced) demonstrated that the evaluation metrics capture distinct aspects of prediction quality, have distinct blind spots, and can vary independently. In the actual DeePEn results, however, models were ranked similarly across metrics, with moderate-to-high correlations between model rankings (SOM Fig. S7). Thus, for model comparison alone, the metrics often supported similar conclusions despite not being interchangeable. However, while rankings across metrics were robust (SOM Fig. S8), the absolute value ranges of the metrics themselves differed substantially. This discrepancy has an important impact on practical usability.

With seemingly high AUCPR scores even at increased depth (Fig. 3, Fig. 4, Table 1), today’s models appear to provide considerable value as prefilter for large-scale high throughput variant screening (SOM Fig. S10). Our observation that Spearman_func_ (Eq. 2), i.e., the Spearman rank correlation was not exceeding 0.4 on the deeper bins (Fig. 3) might caution the pursuit of guided *in silico* evolution [15,35]. When predictive models, such as those examined here, are used to guide generative models toward variants of higher (predicted) fitness, they will mislead the generative models at an increasing rate the further these variants deviate from the wild type. The same applies when using such predictive models to evaluate generative model outputs.

The TOP10% recall values were numerically lower than the Spearman_func_ values that already dropped compared to AUCPR (Fig. 3, Table 1). However, when sampling a few top variants (e.g. a single microwell plate), often the goal rather is to reliably identify a few highly functional variants, rather than all of them. When simulating this scenario of sampling a single 96 well plate per model for DeePEn (Fig. 6), the tested models successfully identified a few of the most effective (top 100) functional variants in three of the four datasets (no model found any of the top *Dehydratase* variants). However, nearly all top functional variants correctly identified were very close (distance 2–3) to the training data. Overall, this analysis once again confirmed our central finding that model predictions are more accurate for shallow variants and deteriorate with increasing depth.

NDCG@10%_func_ (Eq. 4) combines these aspects into a single metric; the relevance score of 0 for non-functional variants severely punishes false positives, implicitly rewarding good binary classification. Like Spearman_func_, NDCG is order dependent. Through the “@k” parameter, it can be focused on the most relevant top model predictions in analogy to the TOP10%_func_ metric. similar to the TOP10%_func_ metric. On the downside, changes in NDCG score are directly linked neither to classification performance, nor to ranking effectiveness, nor to the identification of the top variants. Moreover, these aspects are not weighted equally: for DeePEn, NDCG correlated more with AUCPR (SOM Fig. S7) than the other two metrics. Another major downside of NDCG was the small margin it put between a perfect and a random prediction (Table 1, SOM Table S2). Overall, NDCG@10%_func_ is a useful complementary metric, but its non-linear and relatively unintuitive formulation limits interpretability and argues for its use alongside more transparent metrics.

There are certain limitations to this work we want to point out. First, DeePEn is a retrospective benchmark based on experimentally characterized DMS datasets. It evaluates variant-effect prediction and prioritization on held-out variants, rather than prospective sequence generation, iterative optimization, or experimental success in a goal-directed engineering campaign. DeePEn therefore assesses a predictive component relevant to protein-engineering workflows, not the complete engineering process. Second, we investigated only four datasets. Because they span diverse organisms, phenotypes, assays, and dataset-generation methods, we observed very large dataset-dependent differences in predictive performance (Fig. 4), making it difficult to anticipate how the models would perform on a novel problem. All four datasets were selected to represent data-rich settings, and training deliberately included higher-order variants rather than being restricted to single mutants. Consequently, our findings may not generalize to different circumstances like a low data regime. Third, we retained the binarization thresholds used in ProteinGym. While three out of four were biologically motivated (the fourth, Phototropin, used the median), real engineering campaigns may prefer different thresholds - e.g., producing variants better than wild-type or exceeding a specific design goal. Because these thresholds affect both evaluation across all four metrics and the relevance scores used to train our PT5-LoRA_rank model, we assessed the sensitivity of model performance to the threshold choice. Specifically, we compared normalized AUCPR values for PT5-LoRA_reg using the ProteinGym thresholds and wild-type fitness as alternative cutoffs (SOM Tables S14–S17). In three of the four datasets, identifying variants that outperformed wild type was more challenging than distinguishing functional from non-functional variants; for AAV Capsid, the threshold was already close to wild-type fitness. Finally, our evaluation focuses on relative variant prioritization rather than quantitative calibration, meaning DeePEn does not benchmark predictions against absolute fitness measurements.

## 4. Conclusion

We provided our depth sensitive DeePEn benchmark to assess the ability of models to identify and prioritize experimentally characterized functional variants. Our benchmark revealed several shortcomings in the way models are assessed and relied upon. While we make the benchmark freely available to assist in advancing protein engineering, our focus was largely on raising awareness for a problem which might or might not be solved comprehensively in the future. Future solutions will include better data (most of today’s data is very shallow in terms of mutation depth: Fig. 1), better models (models deteriorate with depth, Fig. 2, 3, 4 & S1), and better metrics (Fig. 5). DeePEn can help colleagues to do the best with what is available; it can assess today’s and future methods. The four metrics included in DeePN are specifically suited to protein engineering and advocate for the use of multi-metric evaluation, as metrics measuring different aspects complement each other. Particularly, relying solely on Spearman’s correlation can easily mislead (Fig. 5).

One fundamental limitation of our approach is that it remains challenging to efficiently sample at higher mutational depth, as experimentally characterized functional variants get increasingly sparse. Either staying close to known experimental data [3] or using active learning approaches , in which prediction models are only used from variants close to existing experimental results and gradually moving further away as more data becomes available round by round, are therefore probably the most reliable options for using current models.

## 5. Methods

### 5.1. Dataset selection

We restricted dataset selection to ProteinGym to retain consistent preprocessing and annotation across datasets; external datasets were not surveyed exhaustively. To arrive at the final four datasets from ProteinGym [1], we followed the following selection process (also see Table S1 for the concrete datasets removed at each step). Out of the 69 DMS datasets from the “Substitution Multiples” subset (sets with mutational depth greater than 1), only 11 had a mutational depth beyond two. Of those 11, three were still too shallow and had no variants above edit distance 4, which was unsuited for the depth focus we were interested in. Three further datasets were excluded because, while being deep enough, they had no functional variants at increasing depth. The last exclusion was performed due to the resulting very small training set size (below 250 datapoints), which did not fit in with the high data setting we wanted to explore.

### 5.2. Datasets

Datasets utilized for our benchmark are shown in Table 2. The raw data were extracted from the ProteinGym dataset and split into training, validation, test set according to the distance from wildtype column in Table 2. A more fine-grained version with statistics for each individual edit distance and each individual evaluation bin is provided in Tables S2-S5.

### 5.3. Model training

We considered two baselines, namely, similarity-based inference (SBI) and one-hot encoding (1-hot) of the input sequences.

For SBI we used MMSeqs2 [39] to search each query (Q) in the test set against all training set variants. The fitness value of the top hit (closest training sequence) determined that for Q. We used the mmseqs map function with default parameters for this, which is specifically set up for the search among very similar sequences.

Biotrainer [40,41] was used to compute the 1-hot baseline. We provided the data files to the end-to-end training pipeline and extracted the final predictions. Biotrainer was configured to utilize one-hot-encoded sequences as input and use a small multi-layer perceptron to predict continuous labels, using a simple mean squared error (MSE) loss.

Zero-shot results were used directly from ProteinGym [1]. ESM-2 used the masked-marginals strategy, masking every mutated position individually. The sum of per-mutation log-odds gives the mutant’s overall fitness score. Contrary to this additive approach, Tranception_L scores were computed using the full bidirectional autoregressive log-likelihood of each mutated sequence (averaged over forward and reverse passes), with inference-time retrieval blending the transformer’s position-wise probabilities with amino-acid frequencies from a homologous MSA.

For MAVE-NN predictions we closely followed the Protein DMS modeling script provided by the authors. We kept the default model and training parameters and tried both, the “additive” and “blackbox” gpmap type, as the latter yielded better results in previous work [9].

The METL predictions were done with both 50M global models, where the 3D model takes the pdb structure of the wildtype protein as additional input. Sequences with a length different from wildtype were again excluded. We created our datasets analog to the “avgfp”-example provided by the authors and finetuned the global 1D and 3D models using the respective example notebook, keeping all parameters with the suggested default settings. Without a Rosetta license we were unable to pretrain protein specific local models for the proteins investigated here.

For our own pLM-based predictions we utilized ProtT5-XL-UniRef50 [16] as the base model and offered four different approaches. The first two were based on pretrained embeddings and training a small, supervised prediction head, a 2-layer MLP with a ReLU-activated hidden layer, dropout regularization, and a final linear projection layer, with the embedding vectors as input, predicting continuous labels, using a simple MSE loss. For the embedding predictor subbed evo we added a short “continued pretraining” phase using a MSA of close homologs to the protein of interest, before extracting embeddings. This approach called evo-tuning was described previously [27].

Following previous research [17], we also applied parameter efficient fine-tuning techniques [42]. We compared a simple MSE regression training to a variety of different training setups (SOM, Table S11) and ended up picking the lambda-ranking loss [32] version as the numerically best performing configuration.

The prediction head was left unchanged to keep results comparable between all pLM predictors. All hyperparameters used for training are provided with the respective training notebooks. For evo-tuning we continued pretraining with the original ProtT5 masking objective (masking 15% of residues in the encoder input_ids). We chose to use LoRA [29] (rank 4) instead of full model training, AdamW [43,44] optimizer with a learning rate of 3e-4. All further details can be found in the evo-tuning notebook in our repository.

For the Baseline model shown in our training depth analysis (Fig. 2 and Fig. S1), we took the 213 remaining (217 - 4 DeePEn sets) DMS sets from ProteinGym, excluding all close homologs to the DeePEn proteins (after computing local alignments we used >20% identity and >35% coverage as thresholds). The resulting 191 we reduced further by removing duplicate sequences (some proteins in ProteinGym have multiple different DMS values for the same sequence) and ended up with 184 datasets. We standardized the DMS scores (per dataset) and used the 1.3 million rows to train PT5-LoRA_rank models (again three random seeds) for one epoch.

### 5.4. Benchmark Evaluation

For the evaluation of our DeePEn benchmark we combined prediction results on all four datasets. Starting with three lists (from different random model seeds) of predictions on each respective test set we first split those into smaller test subsets each covering only a single mutational depth (distance to wildtype). The deepest subset for each dataset covered multiple depths, as the number of sequences at individual depth was often very low as depth increased. We chose the threshold for the last subset such that we always ended up with at least 50 functional sequences in that subset (Tables S2-S5). Next, we calculated all evaluation metrics on those subsets. These results were then grouped into the “shallow (5-9)” and “higher depth (10+)” bins as shown in Table 2. The DeePEn results in Fig.3 and Table 1 show those binned results. To determine the error bars, we did not calculate confidence intervals directly on the binned results because we wanted to avoid measuring differences coming from the underlying data. Instead, we provided a measure of uncertainty only originating from the prediction methods themself. For this we calculated 95% CIs for each individual test subset across the three different random seeds and averaged those CI values across the respective depth bin (and datasets). This also meant not assigning any “error” to deterministic methods like SBI.

### 5.5. Evaluation Metrics

The area under the precision-recall curve (AUCPR) is a single number between 0 and 1 that summarizes the performance of a binary classification model. It represents the area below a curve of precision and recall value pairs for all possible thresholds for a given set of model predictions. A random baseline predictor will result in a score equal to the proportion of positive samples in the dataset. While the widely spread claim, that AUCPR is in general superior to the area under the receiver operating characteristic (AUROC) for imbalanced datasets, is misleading [45], in the specific setting of protein engineering we deliberately wanted to prioritize limiting false positive errors of variants with high predicted scores to avoid wasting experimental budget and therefore chose AUCPR.

As the primary ranking metric, we used Spearman’s rank correlation coefficient. This non-parametric measure assesses the monotonic relationship between two ranked variables, making it suitable for quantifying how closely the predicted ranking matches the ground truth ranking. First, both the predicted and true ranking lists are converted into ranks. Spearman’s rank correlation coefficient is then computed as:

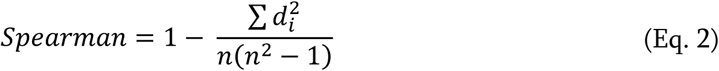

where d_i_ is the difference between the paired ranks of the same item, and n is the number of items ranked. The coefficient ranges from −1 (perfect negative correlation) to +1 (perfect positive correlation), with 0 indicating no correlation. For our experiments we chose to calculate the correlation only for all functional variants within subsets, as those are of interest for protein engineering and dubbed this metric Spearman_func_. First, we filtered out all non-functional variants for each subgroup based on the binary score from ProteinGym and then calculated the Spearman correlation for the remaining sequences based on their DMS score.

To assess the quality of predicted top picks, we employed the Normalized Discounted Cumulative Gain (NDCG) metric. NDCG is widely used in information retrieval to measure the relevance, or gain, of a document based on its position in the result list. The core concept is that highly relevant items appearing lower in the ranking list should be penalized. Given a set of query results with graded relevance, the Discounted Cumulative Gain (DCG) is first calculated as follows:

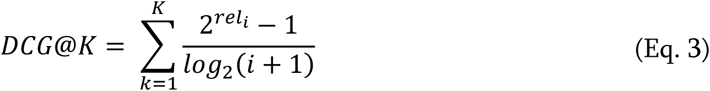

where rel_i_ denotes the relevance score of the item at position i, and the top K positions are taken into consideration for calculating the DCG [32]. For our experiments the relevance score was defined as 0 for all non-functional variants and we used linear min-max scaling of the continuous DMS scores to assign values between 1 (lowest functional score) and 3 (highest functional score) to functional variants. This makes this metric score dependent.

To normalize the score, the Ideal DCG (IDCG) is calculated by sorting the items by their relevance scores and computing DCG for this ideal ranking. NDCG is computed as:

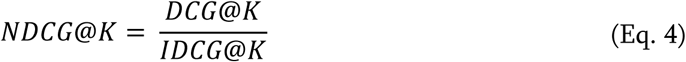

This produces a score between 0 and 1, where 1 represents a perfectly ranked result. Throughout our experiments K was defined to be the top 10% functional variants of each respective test set, and we therefore referred to the metric as NDCG@10%_func_. The score of a random baseline is not obvious for NDCG and was calculated numerically on our datasets.

Lastly, we introduced a simple recall metric which we named TOP10%_func_. We were again only taking the top 10% of functional sequences into account for each respective test set bin. For those n top variants, we checked how many also occurred in the top n predictions and reported this overlapping share. Therefore, if a variant was within the top n predictions the exact ranking did not affect the metric in any way. The score will range from 0 to 1 with a random predictor performance depending on the ratio between functional and non-functional variants. We again calculated the random performance score numerically.

As AUCPR, NDCG@10%_func_ and to a lesser extend TOP10%_func_ are influenced by the prevalence of functional data points in test sets, we normalized those metrics to a range between 0 (random performance) and 1 (perfect) using Eq. 5. Otherwise, comparisons between shallow and higher depth bins and averaging over different depth subsets within a bin would be unevenly influenced by this prevalence. We therefore normalized on a subset level.

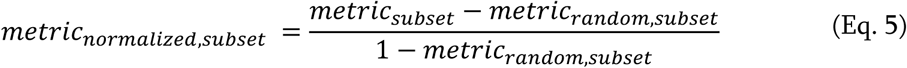

### 5.6. Synthetic dataset for metric independence

Our synthetic dataset consisted of 1 million rows (sequences) with 20% of them put into the “functional” category. NDCG relevance scores were assigned using the same preprocessing stated above. Next, we determined the true ranking of those sequences which can be seen as the “Perfect ranking” in Fig. 4. To make the rankings visually clearer we showed only 100 (every 10.000th row) rows in these ranking plots. The *Random* predictor was created by uniformly shuffling all ranks. For the *Balanced method* (for which we targeted a value of 0.7 for all five evaluation metrics) and the other three *Methods* (#1-3) we manually shuffled and swapped specific parts of the dataset. A detailed reconstruction of all rankings can be found in the “create_display_items_main”-notebook. For *Method #1* we put an entire block of non-functional sequences above the functional ones. The ranking for *Method #2* was created by inverting the ranking of functional variants and increasing the ranking quality of non-functional variants compared to the *Balanced method* baseline, this allowed us to achieve an unchanged overall Spearman of 0.7. For *Method #3* we kept the top 25% of the functional variants well ranked while reducing ranking quality below.

### 5.7. Statistics & Reproducibility

Raw input data, including the four DMS datasets downloaded from ProteinGym, pdb-structures for METL 3D fine-tuning, MSA files for evo-tuning and predictions for all tested models, were made available.

All figures and tables can be reproduced using the corresponding notebooks (one for the main text another for supplemental online materials).

Training code was made available for each of the LoRA fine-tuned ProtT5 variants explored in SOM Table S11. Model checkpoints were made available for our best performing model “PT5-LoRA_rank.” (the lambda_rank model in Table S11). Checkpoints for other models can be provided upon request.

## 6. Data availability

All data used as input, as well as all results used to perform the analysis shown here are available in our repository at https://github.com/RSchmirler/DeePEn. Ready to use data splits are available at https://huggingface.co/datasets/RobSchmi/DeePEn as well. The original raw data is available on https://proteingym.org/ (v1.3). Data for figures and tables is available in the Source Data file.

## 7. Code availability

We made the code to train ProtT5 based prediction models available in our repository (https://github.com/RSchmirler/DeePEn), together with notebooks to reproduce all analysis and display items. Training code for METL [23] local models is available in their respective repository (https://github.com/gitter-lab/metl, version 0.1.0). For MAVE-NN [24]we used the mavenn python package version 1.1.3. The one-hot-encoding baseline model utilized biotrainer which is now part of Biocentral [41] (available at https://github.com/biocentral/biocentral, version 1.3.0). For the similarity-based-inference baseline we used MMseqs2 [39] (https://github.com/soedinglab/MMseqs2, version 14.7e284).

## 9. Acknowledgments

We thank Nikita Kugut (TUM) for his support with many aspects of this work. We thank the Technical University of Munich (TUM) for providing facilities and resources. Finally, we thank those who deposit experimental data in public databases, maintain these databases, and make their resources publicly available.

## Author Contributions Statement

R.S., M.H. and B.R. contributed to the conception of this study. M.B. provided code and consultation for the “PT5-LoRA Rank. lambda_rank” model and Fig. 4 visualization. S.F. computed the one-hot-encoding baseline utilizing biotrainer. R.S. performed all other modelling work, results analysis, and creation of display items. The first draft of the manuscript was written by R.S.. All authors commented on and refined subsequent versions of the manuscript.

