## Supplementary Online Material for "DeePEn-A Depth sensitive benchmark for Protein Engineering"

### Supplementary Online Material
### for
### Fine-DeePEn-A Depth sensitive benchmark for Protein Engineering

**Robert Schmirler [1,2*], Maurice Brenner [1,3], Sebastian Franz [1], Michael Heinzinger [1,3] & Burkhard Rost [1,4]**

1 TUM (Technical University of Munich), School of Computation, Information and Technology (CIT), Faculty of Informatics, Chair of Bioinformatics & Computational Biology - i12, Boltzmannstr. 3, 85748 Garching/Munich, Germany
2 AbbVie Deutschland GmbH & Co. KG, Innovation Center, BTS IR LU, Ludwigshafen, 67061, Germany
3 Institute of Computational Biology, Computational Health Center, Helmholtz Munich, Munich, 85764, Germany
4 Institute for Advanced Study (TUM-IAS), Lichtenbergstr. 2a, 85748 Garching/Munich, Germany & TUM School of Life Sciences Weihenstephan (WZW), Alte Akademie 8, Freising, Germany

### 1. SOM 1: ProteinGym dataset prefiltering

Here we provide a detailed step by step analysis of which datasets we excluded and the respective exclusion criteria. We start with all 69 datasets which contain higher order mutations, this subset can be downloaded as „Substitution Multiples“ from the ProteinGym download webpage.

**Table S1: ProteinGym dataset prefiltering**

| Number of DMS datasets before | Filter criteria | Datasets removed | Number of DMS datasets removed |
|---|---|---|---|
| 69 („Substitution Multiples“ subset) | Removed datasets only containing double mutants as their highest depth | 50x Tsuboyama et al.'s Mega-scale experiment<br>A4_HUMAN_Seuma_2022<br>DLG4_HUMAN_Faure_2021<br>GRB2_HUMAN_Faure_2021<br>PABP_YEAST_Melamed_2013<br>RASK_HUMAN_Weng_2022_abundance<br>RASK_HUMAN_Weng_2022_[…]_K55<br>SPG1_STRSG_Olson_2014<br>YAP1_HUMAN_Araya_2012 | 58 |
| 11 | Max depth <5, which would result in no test set | F7YBW7_MESOW_Ding_2023<br>F7YBW8_MESOW_Aakre_2015<br>SPG1_STRSG_Wu_2016 | 3 |
| 8 | No functional variants at higher depth (10+) | Q6WV13_9MAXI_Somermeyer_2022<br>Q8WTC7_9CNID_Somermeyer_2022<br>D7PM05_CLYGR_Somermeyer_2022 | 3 |
| 5 | Small data set size | GCN4_YEAST_Staller_2018 | 1 |
| Selected datasets: | | PHOT_CHLRE_Chen_2023 (Phototropin)<br>HIS7_YEAST_Pokusaeva_2019 (Dehydratase)<br>GFP_AEQVI_Sarkisyan_2016 (GFP)<br>CAPSD_AAV2S_Sinai_2021 (AAV capsid) | |

### 2. SOM 2: DeePEn results -absolute values

---

**Table S2: DeePEn results - absolute metric values** Δ

| *Model* | *Depth* | *AUCPR* | *Spearman$_{func}$* | *TOP10%$_{func}$* | *NDCG@10%$_{func}$* |
|---|---|---|---|---|---|
| *Random baseline* | *shallow (5-9)* | 49 ± 0 | 0 ± 0 | 5 ± 0 | 38 ± 0 |
| | *higher depth (10+)* | 30 ± 0 | 0 ± 0 | 3 ± 0 | 24 ± 0 |
| *Perfect classifier baseline* | *shallow (5-9)* | 100 ± 0 | 0 ± 0 | 10 ± 0 | 77 ± 0 |
| | *higher depth (10+)* | 100 ± 0 | 0 ± 0 | 10 ± 0 | 77 ± 0 |
| *SBI baseline* | *shallow (5-9)* | 66 ± 0 | 21 ± 0 | 15 ± 0 | 59 ± 0 |
| | *higher depth (10+)* | 39 ± 0 | 12 ± 0 | 6 ± 0 | 34 ± 0 |
| *1-hot baseline* | *shallow (5-9)* | 55 ± 9 | 2 ± 12 | 6 ± 5 | 41 ± 19 |
| | *higher depth (10+)* | 36 ± 10 | 4 ± 19 | 4 ± 4 | 23 ± 22 |
| *ESM2_650M$_{zero-shot}$* | *shallow (5-9)* | 65 ± 0 | 22 ± 0 | 17 ± 0 | 64 ± 0 |
| | *higher depth (10+)* | 61 ± 0 | 34 ± 0 | 20 ± 0 | 65 ± 0 |
| *Tranception_L$_{zero-shot}$* | *shallow (5-9)* | 75 ± 0 | 27 ± 0 | 17 ± 0 | 77 ± 0 |
| | *higher depth (10+)* | 57 ± 0 | 30 ± 0 | 18 ± 0 | 70 ± 0 |
| *MAVE-NN_add* | *shallow (5-9)* | 87 ± 1 | 42 ± 2 | 28 ± 2 | 86 ± 1 |
| | *higher depth (10+)* | 56 ± 1 | 25 ± 1 | 25 ± 2 | 64 ± 2 |
| *MAVE-NN_bb* | *shallow (5-9)* | 87 ± 6 | 45 ± 18 | 31 ± 11 | 82 ± 10 |
| | *higher depth (10+)* | 62 ± 8 | 28 ± 19 | 22 ± 7 | 59 ± 9 |
| *METL-global_1D* | *shallow (5-9)* | 87 ± 1 | 47 ± 3 | 32 ± 4 | 84 ± 4 |
| | *higher depth (10+)* | 66 ± 3 | 31 ± 7 | 22 ± 6 | 63 ± 7 |
| *METL-global_3D* | *shallow (5-9)* | 85 ± 2 | 48 ± 3 | 33 ± 3 | 83 ± 4 |
| | *higher depth (10+)* | 69 ± 4 | 35 ± 6 | 27 ± 6 | 70 ± 6 |
| *PT5-LoRA_reg* | *shallow (5-9)* | 90 ± 3 | 53 ± 4 | 37 ± 6 | 85 ± 7 |
| | *higher depth (10+)* | 75 ± 11 | 36 ± 8 | 28 ± 13 | 70 ± 19 |
| *PT5-LoRA_rank* | *shallow (5-9)* | 90 ± 1 | 54 ± 4 | 36 ± 7 | 89 ± 2 |
| | *higher depth (10+)* | 76 ± 8 | 39 ± 13 | 30 ± 10 | 78 ± 15 |
| *PT5-Emb* | *shallow (5-9)* | 86 ± 2 | 46 ± 5 | 28 ± 5 | 83 ± 4 |
| | *higher depth (10+)* | 69 ± 6 | 35 ± 9 | 21 ± 12 | 66 ± 20 |
| *PT5-Emb_evo* | *shallow (5-9)* | 90 ± 2 | 49 ± 4 | 33 ± 6 | 86 ± 5 |
| | *higher depth (10+)* | 74 ± 8 | 32 ± 20 | 23 ± 16 | 75 ± 20 |

Δ Results per model for both depth bins (shallow/higher) give averages over all depth subsets and all four datasets. 95% confidence intervals of the mean values were only calculated for model replicates within individual depths/dataset-combinations. Metric values multiplied by 100, numerically best performing model per depth and metric in bold, model differences are not statistically significant. The Random baseline is randomly shuffling all variants, while the Perfect classifier baseline is perfectly separating non-functional and functional variants while randomly shuffling all functional variants.

### 3. SOM 3: DeePEn dataset subsets

This section provides detailed information about the respective evaluation subsets. For the DeePEn benchmark metric calculation (Table 1, Fig. 3) each row of the test split (with the respective distance from wildtype) is handled as an individual test subset during metric calculation. The "shallow" and "higher depth" bin assignment and averaging is done based on the distance bins shown here.

**Table S3: AAV capsid - detailed**

| Split | Distance from wildtype | Distance bin | Number of seq | Number of functional seq | Share of functional seq |
|---|---|---|---|---|---|
| Training | 1 | 1-3 | 532 | 315 | 59.2 % |
| | 2 | | 10,833 | 4,601 | 42.5 % |
| | 3 | | 6,906 | 5,177 | 75.0 % |
| Validation | 4 | 4 | 6,646 | 4,504 | 67.8 % |
| Test | 5 | 5-9 | 6,298 | 3,772 | 59.9 % |
| | 6 | | 5,396 | 2,584 | 47.9 % |
| | 7 | | 858 | 398 | 46.4 % |
| | 8 | | 739 | 237 | 32.1 % |
| | 9 | | 651 | 141 | 21.7 % |
| | 10 | 10-28 | 583 | 92 | 15.8 % |
| | 11-28 | | 2,886 | 91 | 3.2 % |

**Table S4: Dehydratase - detailed**

| Split | Distance from wildtype | Distance bin | Number of seq | Number of functional seq | Share of functional seq |
|---|---|---|---|---|---|
| Training | 1 | 1-3 | 168 | 162 | 96.4 % |
| | 2 | | 1,475 | 1,375 | 93.2 % |
| | 3 | | 7,627 | 6,661 | 87.3 % |
| Validation | 4 | 4 | 25,927 | 21,177 | 81.7 % |
| Test | 5 | 5-9 | 60,263 | 45,987 | 76.3 % |
| | 6 | | 98,869 | 70,269 | 71.1 % |
| | 7 | | 115,751 | 76,096 | 65.7 % |
| | 8 | | 97,639 | 58,953 | 60.4 % |
| | 9 | | 57,139 | 32,216 | 56.4 % |
| | 10 | 10-28 | 22,942 | 12,488 | 54.4 % |
| | 11 | | 6,370 | 3,727 | 58.5 % |
| | 12 | | 1,354 | 920 | 67.9 % |
| | 13-28 | | 613 | 173 | 28.2 % |

#### Table S5: GFP - detailed

| Split | Distance from wildtype | Distance bin | Number of seq | Number of functional seq | Share of functional seq |
|---|---|---|---|---|---|
| Training | 1 | 1-3 | 1,084 | 993 | 91.6 % |
| | 2 | | 12,777 | 11,367 | 89.0 % |
| | 3 | | 12,336 | 9,234 | 74.9 % |
| Validation | 4 | 4 | 9,387 | 5,104 | 54.4 % |
| Test | 5 | 5-7 | 6,825 | 2,284 | 33.5 % |
| | 6 | | 4,298 | 802 | 18.7 % |
| | 7 | | 2,526 | 229 | 9.1 % |
| | 8-15*: 8* | 8-15 | 1,364 | 61 | 4.5 % |
| | 8-15*: 9* | | 627 | 10 | 1.6 % |
| | 8-15*: 10-15* | | 490 | 2 | 0.4 % |

* we wanted to show the detailed distribution of functional sequences for the 8-15 evaluation subset and therefore showed individual values for depth 8, 9 and 10-15 here. Metric calculation was done over the entire 8-15 subset, which means the *GFP* dataset only contributes a single subset to the “higher depth (10+)” bin.

#### Table S6: Phototropin - detailed

| Split | Distance from wildtype | Distance bin | Number of seq | Number of functional seq | Share of functional seq |
|---|---|---|---|---|---|
| Training | 1 | 1-3 | 2,122 | 1,828 | 86.1 % |
| | 2 | | 176 | 168 | 95.5 % |
| | 3 | | 978 | 856 | 87.5 % |
| Validation | 4 | 4 | 3,565 | 2,859 | 80.2 % |
| Test | 5 | 5-9 | 9,603 | 7,015 | 73.1 % |
| | 6 | | 19,193 | 12,518 | 65.2 % |
| | 7 | | 29,255 | 16,938 | 57.9 % |
| | 8 | | 34,230 | 17,071 | 49.9 % |
| | 9 | | 30,925 | 13,010 | 42.1 % |
| | 10 | 10-15 | 21,167 | 7,361 | 34.8 % |
| | 11 | | 11,022 | 3,057 | 27.7 % |
| | 12 | | 4,093 | 905 | 22.1 % |
| | 13-15 | | 1,200 | 182 | 15.2 % |

### 4. SOM 4: Training depth effect

Here we show the analog results to Fig. 2 for the other three datasets. The training depth 1-3, which we chose for the DeePEn benchmark, yielded high prediction performance across datasets with remaining room to improve. The exception was the *GFP* dataset, where high prediction performance for higher depth (depth 7 and 8-15) remained elusive even when training on all shallower data (training depth 1-6).

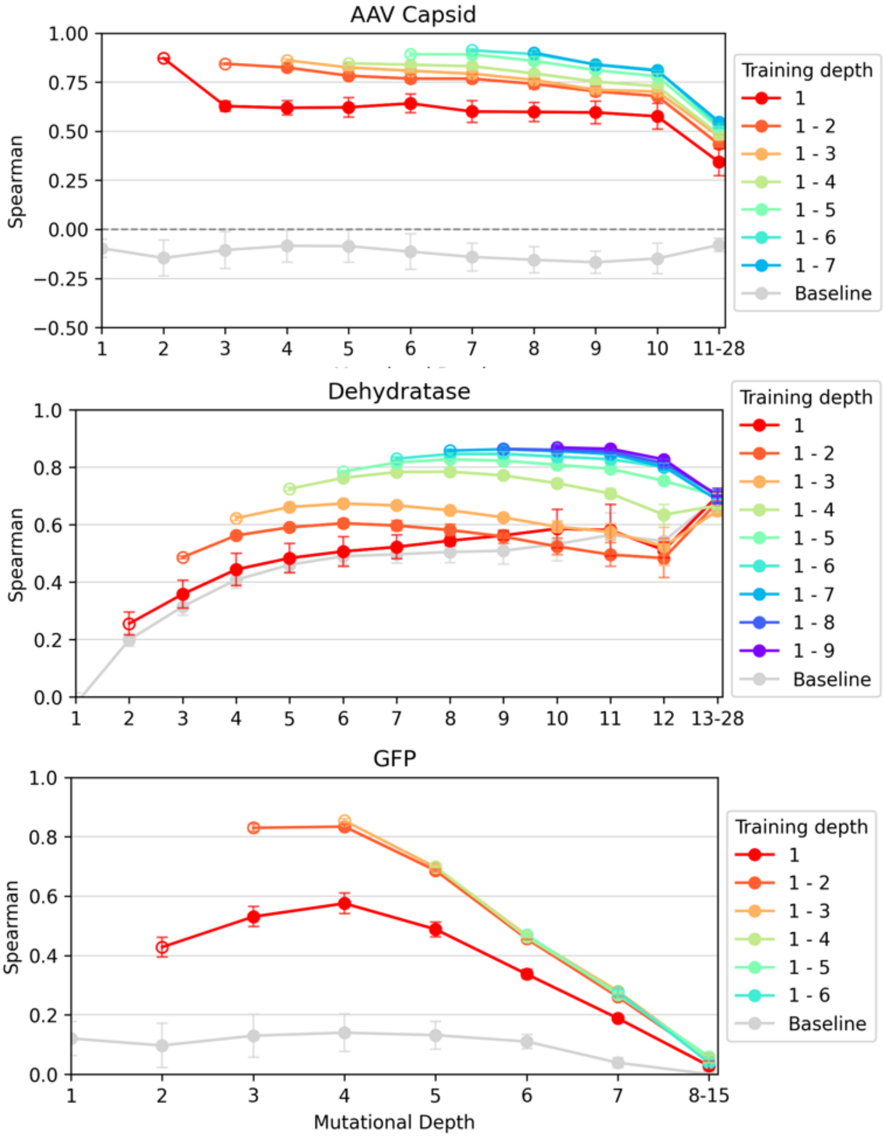


**Fig. S1: Effect of increasing training depth.** Lines are colored by the depth used for training (1-3 means that training contained mutations with d<4). Each point represents the Spearman correlation between predicted and actual values for a specific mutational depth. The leftmost point (for training depth 1-3 the validation is d=4) of each line was used for validation (empty circle), while all other points were the test set (full circles). Different training depth significantly impacted model capability. Across the three datasets shown here only utilizing SAVs was not sufficient to achieve high predictive performance (red lines with training depth 1). When adding doubles to the training data (red lines, training depth 1-2) performance was much better across all test depths. Further adding triplets, did also increase performance (orange lines with training depth 1-3) for two (*AAV Capsid* and *Dehydratase*) out of these three datasets. The Baseline was trained on 184 non-homolog ProteinGym DMS sets.

### 5. SOM 5: DeePEn results for individual datasets

Instead of aggregating over all datasets, we provided results for the four individual datasets here. Tables S7-S10 (for table caption see Table S8) are analog to Table 1 of the main manuscript. Fig. S2-S5 (for figure caption see Fig. S2) are produced analog to Fig. 3. of the main manuscript. Here the boxplots only showed the distribution within a depth bin for the specific dataset. When a boxplot shows a wide spread, this variance can either originate from model randomness (large CI95 intervals) or from large subset differences inside a bin (tight CI95 intervals). We only provide normalized metric values here (as well as the absolute random baseline values in the first row), complete results tables with absolute values can be found in the repository under *results/main/random_baseline.*

**Table S7: AAV capsid results**$^{\Delta}$

| *Model* | *Depth* | *AUCPR* | *Spearman$_{func}$* | *TOP10%$_{func}$* | *NDCG@10%$_{func}$* |
|---|---|---|---|---|---|
| *Random baseline* | *shallow (5-9)* | 42 ± 0 | 0 ± 0 | 4 ± 0 | 30 ± 0 |
| | *higher depth (10+)* | 10 ± 0 | 0 ± 1 | 1 ± 0 | 7 ± 1 |

| *Model* | *Depth* | *AUCPR norm* | *Spearman$_{func}$* | *TOP10%$_{func}$ norm* | *NDCG@10%$_{func}$ norm* |
|---|---|---|---|---|---|
| *Perfect classifier baseline* | *shallow (5-9)* | 100 ± 0 | 0 ± 0 | 6 ± 0 | 59 ± 0 |
| | *higher depth (10+)* | 100 ± 0 | 0 ± 1 | 9 ± 1 | 73 ± 0 |
| *SBI baseline* | *shallow (5-9)* | 37 ± 0 | 19 ± 0 | 11 ± 0 | 30 ± 0 |
| | *higher depth (10+)* | 9 ± 0 | 3 ± 0 | -1 ± 0 | 12 ± 0 |
| *1-hot baseline* | *shallow (5-9)* | 12 ± 14 | -4 ± 14 | 0 ± 6 | 6 ± 20 |
| | *higher depth (10+)* | 3 ± 6 | -1 ± 18 | -1 ± 0 | -3 ± 11 |
| *ESM2_650M$_{zero-shot}$* | *shallow (5-9)* | 15 ± 0 | 6 ± 0 | 6 ± 0 | 23 ± 0 |
| | *higher depth (10+)* | 4 ± 0 | 15 ± 0 | 5 ± 0 | 12 ± 0 |
| *Tranception_L$_{zero-shot}$* | *shallow (5-9)* | 40 ± 0 | 24 ± 0 | 12 ± 0 | 51 ± 0 |
| | *higher depth (10+)* | 17 ± 0 | 24 ± 0 | 16 ± 0 | 47 ± 0 |
| *MAVE-NN_add* | *shallow (5-9)* | 65 ± 1 | 28 ± 3 | 12 ± 3 | 56 ± 2 |
| | *higher depth (10+)* | 19 ± 1 | 15 ± 2 | 16 ± 0 | 33 ± 7 |
| *MAVE-NN_bb* | *shallow (5-9)* | 52 ± 30 | 32 ± 45 | 18 ± 31 | 45 ± 48 |
| | *higher depth (10+)* | 12 ± 14 | 22 ± 34 | 5 ± 14 | 11 ± 27 |
| *METL-global_1D* | *shallow (5-9)* | 56 ± 2 | 47 ± 3 | 31 ± 7 | 55 ± 12 |
| | *higher depth (10+)* | 25 ± 4 | 30 ± 6 | 12 ± 24 | 23 ± 15 |
| *METL-global_3D* | *shallow (5-9)* | 45 ± 2 | 48 ± 2 | 34 ± 5 | 53 ± 5 |
| | *higher depth (10+)* | 16 ± 1 | **41 ± 4** | 8 ± 8 | 11 ± 7 |
| *PT5-LoRA_reg* | *shallow (5-9)* | 64 ± 7 | **53 ± 5** | **37 ± 9** | 62 ± 18 |
| | *higher depth (10+)* | 26 ± 12 | 38 ± 9 | 22 ± 14 | 31 ± 39 |
| *PT5-LoRA_rank* | *shallow (5-9)* | 65 ± 2 | **53 ± 4** | **37 ± 9** | **71 ± 5** |
| | *higher depth (10+)* | 26 ± 4 | **41 ± 6** | **23 ± 8** | 46 ± 20 |
| *PT5-Emb* | *shallow (5-9)* | 64 ± 5 | 46 ± 6 | 28 ± 6 | 67 ± 7 |
| | *higher depth (10+)* | 21 ± 2 | 27 ± 7 | 7 ± 8 | 17 ± 19 |
| *PT5-Emb_evo* | *shallow (5-9)* | **67 ± 8** | 50 ± 4 | 32 ± 11 | 67 ± 15 |
| | *higher depth (10+)* | **31 ± 7** | 25 ± 12 | 7 ± 21 | **47 ± 43** |

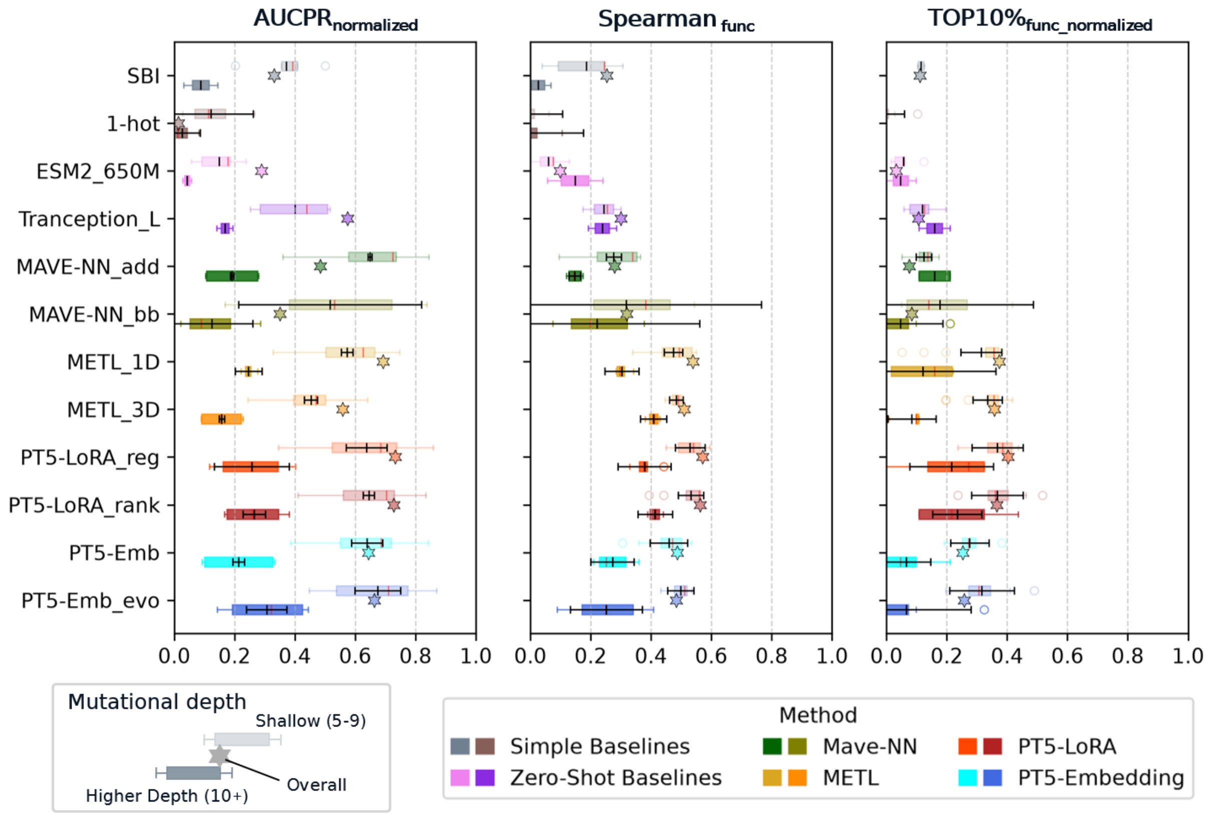


**Fig. S2: AAV capsid results** †

† Methods: in gray simple baselines: SIB - Similarity-based inference and 1-hot - as one-hot encoding (training on amino acid sequence in one-hot encoding). Zero-shot baselines in pink: ESM2_650M (additive masked log-odds) and Tranception_L (autoregressive joint likelihood). MAVE-NN_add uses a fixed additive effect model while the black box variant trains a neural network for effect aggregation. PT5-LoRA models are LoRA finetuned ProtT5 models, while PT5-Emb utilizes frozen ProtT5 embeddings. METL-3D is structure informed while 1D is not. Two boxplots per model show the distribution of the respective metric across depth subsets (within the shallow/higher bins). 95% confidence intervals of the mean values were only calculated for model replicates within individual depths subsets, which allowed capturing only uncertainties originating from the models. To control for shifting class prevalence, metrics are normalized (per subset) to a range between 0 (random) and 1 (perfect prediction).

#### Table S8: Dehydratase results$^{\Delta}$

| *Model* | *Depth* | *AUCPR* | *Spearman$_{func}$* | *TOP10%$_{func}$* | *NDCG@10%$_{func}$* |
|---|---|---|---|---|---|
| *Random baseline* | *shallow (5-9)* | 66 ± 0 | 0 ± 0 | 7 ± 0 | 59 ± 0 |
| | *higher depth (10+)* | 53 ± 0 | 0 ± 0 | 5 ± 0 | 46 ± 0 |
| *Model* | *Depth* | *AUCPR norm* | *Spearman$_{func}$* | *TOP10%$_{func}$ norm* | *NDCG@10%$_{func}$ norm* |
| *Perfect classifier baseline* | *shallow (5-9)* | 100 ± 0 | 0 ± 0 | 4 ± 0 | 73 ± 0 |
| | *higher depth (10+)* | 100 ± 0 | 0 ± 0 | 5 ± 0 | 75 ± 0 |
| *SBI baseline* | *shallow (5-9)* | 51 ± 0 | 10 ± 0 | 6 ± 0 | 42 ± 0 |
| | *higher depth (10+)* | 28 ± 0 | 3 ± 0 | 3 ± 0 | 22 ± 0 |
| *1-hot baseline* | *shallow (5-9)* | 32 ± 7 | 3 ± 4 | 3 ± 1 | 9 ± 18 |
| | *higher depth (10+)* | 23 ± 7 | 6 ± 17 | 0 ± 3 | -15 ± 22 |
| *ESM2_650M$_{zero\text{-}shot}$* | *shallow (5-9)* | 49 ± 0 | 21 ± 0 | **7 ± 0** | 76 ± 0 |
| | *higher depth (10+)* | 83 ± 0 | 40 ± 0 | 12 ± 0 | **81 ± 0** |
| *Tranception_L$_{zero\text{-}shot}$* | *shallow (5-9)* | 77 ± 0 | 27 ± 0 | 4 ± 0 | 73 ± 0 |
| | *higher depth (10+)* | **84 ± 0** | **42 ± 0** | **16 ± 0** | 77 ± 0 |
| *MAVE-NN_add* | *shallow (5-9)* | 65 ± 3 | 9 ± 2 | 0 ± 1 | 76 ± 2 |
| | *higher depth (10+)* | -7 ± 5 | -9 ± 2 | -4 ± 0 | 10 ± 4 |
| *MAVE-NN_bb* | *shallow (5-9)* | 73 ± 5 | 18 ± 14 | 2 ± 4 | 63 ± 2 |
| | *higher depth (10+)* | 12 ± 16 | -3 ± 22 | -3 ± 1 | -2 ± 11 |
| *METL-global_1D* | *shallow (5-9)* | 73 ± 3 | 17 ± 3 | -1 ± 1 | 68 ± 6 |
| | *higher depth (10+)* | 43 ± 6 | 4 ± 13 | -3 ± 1 | 14 ± 17 |
| *METL-global_3D* | *shallow (5-9)* | 76 ± 2 | 24 ± 2 | 2 ± 1 | 67 ± 6 |
| | *higher depth (10+)* | 61 ± 12 | 8 ± 8 | 8 ± 8 | 50 ± 19 |
| *PT5-LoRA_reg* | *shallow (5-9)* | 77 ± 7 | 26 ± 4 | 6 ± 3 | 65 ± 15 |
| | *higher depth (10+)* | 65 ± 17 | 8 ± 4 | 3 ± 10 | 35 ± 37 |
| *PT5-LoRA_rank* | *shallow (5-9)* | 83 ± 6 | 28 ± 5 | 3 ± 7 | **77 ± 7** |
| | *higher depth (10+)* | 69 ± 30 | 10 ± 22 | 4 ± 11 | 56 ± 53 |
| *PT5-Emb* | *shallow (5-9)* | 87 ± 3 | 28 ± 4 | 4 ± 3 | 75 ± 5 |
| | *higher depth (10+)* | 75 ± 19 | 16 ± 11 | 2 ± 9 | 54 ± 40 |
| *PT5-Emb_evo* | *shallow (5-9)* | **88 ± 2** | **30 ± 6** | 5 ± 4 | 73 ± 8 |
| | *higher depth (10+)* | 78 ± 13 | 7 ± 33 | 0 ± 8 | 52 ± 47 |

$^{\Delta}$ Results per model for both depth bins (shallow/higher) give averages over all depth subsets and all four datasets. Baseline methods on top, supervised advanced methods below. To control for shifting class prevalence, metrics are normalized (per depth subset) to a range between 0 (random) and 1 (perfect prediction). Spearman$_{func}$ is not affected as its random performance is already 0. We provide absolute values for the Random baseline in the first row, for all other methods absolute values can be found in the repository. The *Random baseline* is randomly ranking all variants, while the *Perfect classifier baseline* is perfectly separating non-functional (at the bottom) and functional variants (at the top) while randomly shuffling all functional variants. 95% confidence intervals of the mean values were only calculated for model replicates within individual depths subsets. Metric values multiplied by 100, numerically best performing model per depth and metric in bold, these model differences are not statistically significant.

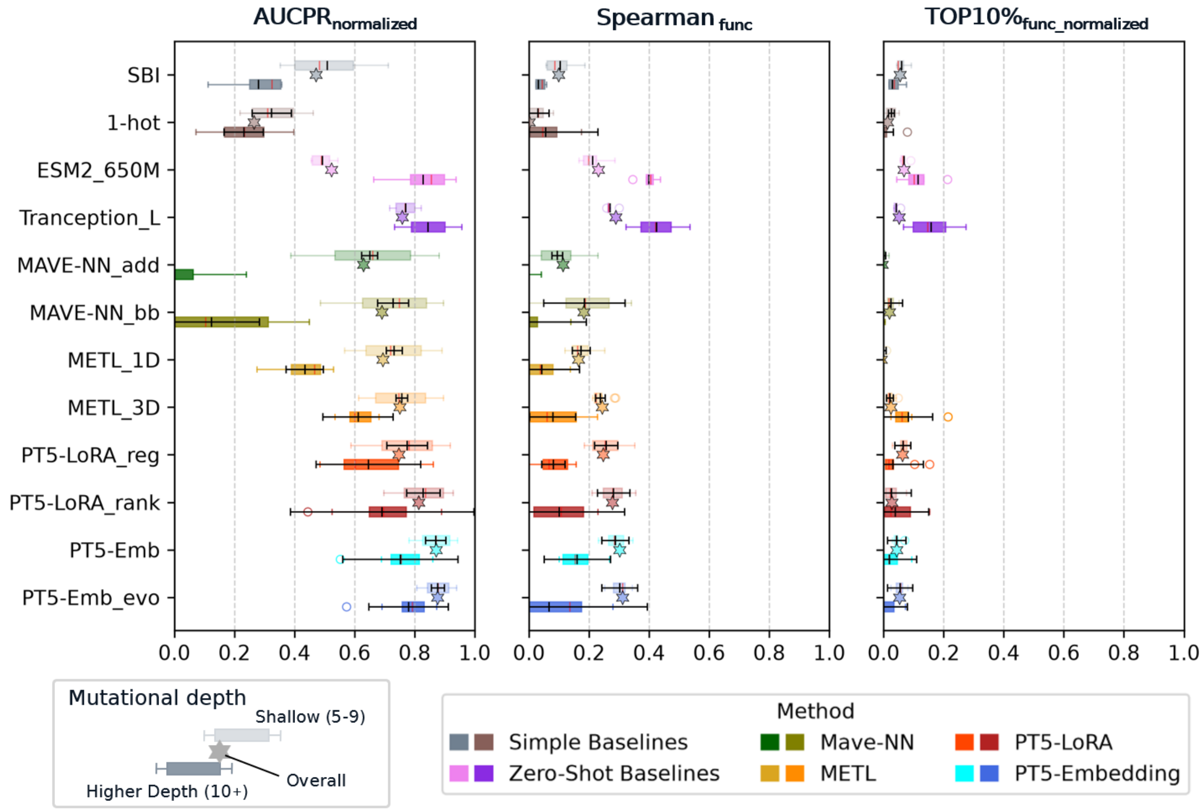


**Fig. S3: Dehydratase results** †

#### Table S9: GFP results$^{\Delta}$

| *Model* | *Depth* | *Spearman* | *AUCPR* | *Spearman*$_{func}$ | *TOP10%*$_{func}$ | *NDCG@10%*$_{func}$ |
|---|---|---|---|---|---|---|
| *Random baseline* | *shallow (5-9)* | 0 ± 0 | 21 ± 0 | 0 ± 0 | 2 ± 0 | 17 ± 0 |
| | *higher depth (10+)* | 0 ± 0 | 3 ± 0 | -1 ± 1 | 0 ± 0 | 3 ± 0 |
| *Model* | *Depth* | *Spearman* | *AUCPR* *norm* | *Spearman*$_{func}$ | *TOP10%*$_{func}$ *norm* | *NDCG@10%*$_{func}$ *norm* |
| *Perfect classifier baseline* | *shallow (5-7)* | 65 ± 0 | 100 ± 0 | 0 ± 0 | 8 ± 0 | 80 ± 0 |
| | *higher depth (8-15)* | 28 ± 0 | 100 ± 0 | -1 ± 1 | 9 ± 1 | 82 ± 0 |
| *SBI baseline* | *shallow (5-7)* | 27 ± 0 | 22 ± 0 | 17 ± 0 | 5 ± 0 | 28 ± 0 |
| | *higher depth (8-15)* | 2 ± 0 | 5 ± 0 | 28 ± 0 | 0 ± 0 | 10 ± 0 |
| *1-hot baseline* | *shallow (5-7)* | -2 ± 13 | -1 ± 8 | -1 ± 11 | -1 ± 1 | -4 ± 28 |
| | *higher depth (8-15)* | -1 ± 2 | 0 ± 1 | -8 ± 1 | 0 ± 0 | -3 ± 0 |
| *ESM2_650M*$_{zero\text{-}shot}$ | *shallow (5-7)* | 1 ± 0 | 1 ± 0 | -3 ± 0 | -1 ± 0 | -6 ± 0 |
| | *higher depth (8-15)* | 2 ± 0 | 1 ± 0 | 1 ± 0 | 0 ± 0 | -3 ± 0 |
| *Tranception_L*$_{zero\text{-}shot}$ | *shallow (5-7)* | 31 ± 0 | 46 ± 0 | 17 ± 0 | 8 ± 0 | 61 ± 0 |
| | *higher depth (8-15)* | 5 ± 0 | 23 ± 0 | 31 ± 0 | 14 ± 0 | 71 ± 0 |
| *MAVE-NN_add* | *shallow (5-7)* | 48 ± 1 | **96 ± 1** | **66 ± 1** | 39 ± 4 | **94 ± 1** |
| | *higher depth (8-15)* | 6 ± 1 | **85 ± 2** | **55 ± 3** | 52 ± 20 | **97 ± 1** |
| *MAVE-NN_bb* | *shallow (5-7)* | **49 ± 10** | **96 ± 2** | 64 ± 2 | **40 ± 8** | **94 ± 1** |
| | *higher depth (8-15)* | 8 ± 9 | **85 ± 2** | 53 ± 7 | **56 ± 0** | **97 ± 1** |
| *METL-global_1D* | *shallow (5-7)* | 47 ± 1 | 94 ± 1 | 52 ± 3 | 30 ± 10 | 93 ± 2 |
| | *higher depth (8-15)* | **9 ± 2** | 80 ± 2 | 35 ± 14 | 28 ± 0 | 92 ± 2 |
| *METL-global_3D* | *shallow (5-7)* | 46 ± 1 | 89 ± 9 | 40 ± 12 | 21 ± 7 | 83 ± 14 |
| | *higher depth (8-15)* | 8 ± 2 | 70 ± 11 | 33 ± 8 | 28 ± 0 | 88 ± 3 |
| *PT5-LoRA_reg* | *shallow (5-7)* | 48 ± 1 | 94 ± 2 | 54 ± 4 | 30 ± 9 | 93 ± 1 |
| | *higher depth (8-15)* | 6 ± 3 | 81 ± 5 | 45 ± 15 | 38 ± 20 | 95 ± 3 |
| *PT5-LoRA_rank* | *shallow (5-7)* | 47 ± 1 | 94 ± 2 | 56 ± 4 | 32 ± 13 | 93 ± 2 |
| | *higher depth (8-15)* | 4 ± 1 | 80 ± 3 | 44 ± 7 | 47 ± 20 | 96 ± 1 |
| *PT5-Emb* | *shallow (5-7)* | 43 ± 1 | 65 ± 4 | 23 ± 7 | 11 ± 7 | 63 ± 13 |
| | *higher depth (8-15)* | 3 ± 3 | 22 ± 6 | 28 ± 10 | 4 ± 20 | 28 ± 52 |
| *PT5-Emb_evo* | *shallow (5-7)* | 47 ± 1 | 87 ± 2 | 34 ± 3 | 18 ± 9 | 85 ± 5 |
| | *higher depth (8-15)* | 5 ± 2 | 56 ± 12 | 36 ± 17 | 28 ± 36 | 70 ± 4 |

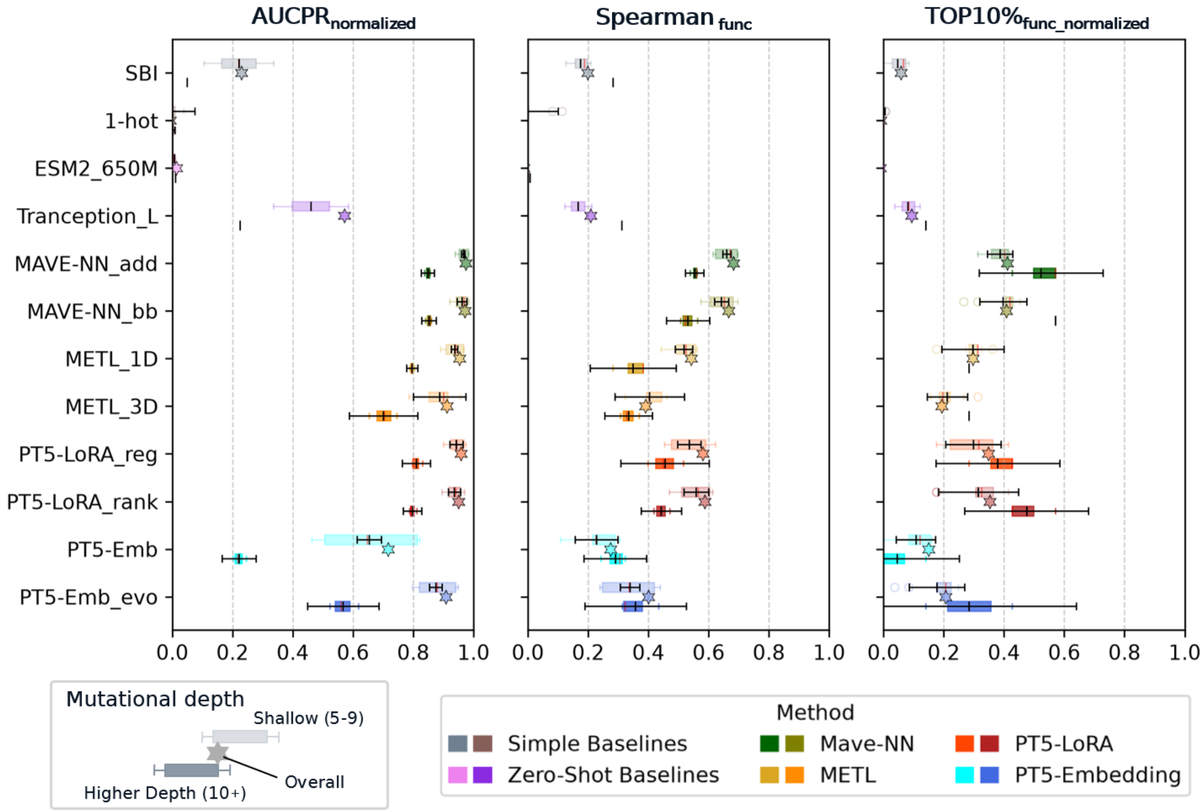


**Fig. S4: GFP results †**

#### Table S10: Phototropin results[Δ]

| *Model* | *Depth* | *AUCPR* | *Spearman$_{func}$* | *TOP10%$_{func}$* | *NDCG@10%$_{func}$* |
|---|---|---|---|---|---|
| *Random baseline* | *shallow (5-9)* | 57 ± 0 | 0 ± 0 | 6 ± 0 | 39 ± 0 |
| | *higher depth (10+)* | 25 ± 0 | 0 ± 0 | 3 ± 0 | 16 ± 0 |
| *Model* | *Depth* | *AUCPR norm* | *Spearman$_{func}$* | *TOP10%$_{func}$ norm* | *NDCG@10%$_{func}$ norm* |
| *Perfect classifier baseline* | *shallow (5-9)* | 100 ± 0 | 0 ± 0 | 4 ± 0 | 46 ± 0 |
| | *higher depth (10+)* | 100 ± 0 | 0 ± 0 | 7 ± 0 | 59 ± 0 |
| *SBI baseline* | *shallow (5-9)* | 34 ± 0 | 36 ± 0 | 17 ± 0 | 41 ± 0 |
| | *higher depth (10+)* | 8 ± 0 | 22 ± 0 | 5 ± 0 | 10 ± 0 |
| *1-hot baseline* | *shallow (5-9)* | 11 ± 44 | 7 ± 19 | 2 ± 11 | 11 ± 55 |
| | *higher depth (10+)* | 7 ± 28 | 9 ± 25 | 2 ± 10 | 9 ± 53 |
| *ESM2_650M$_{zero-shot}$* | *shallow (5-9)* | 76 ± 0 | 53 ± 0 | 34 ± 0 | 77 ± 0 |
| | *higher depth (10+)* | 57 ± 0 | 47 ± 0 | 34 ± 0 | 74 ± 0 |
| *Tranception_L$_{zero-shot}$* | *shallow (5-9)* | 60 ± 0 | 35 ± 0 | 25 ± 0 | 69 ± 0 |
| | *higher depth (10+)* | 33 ± 0 | 20 ± 0 | 14 ± 0 | 54 ± 0 |
| *MAVE-NN_add* | *shallow (5-9)* | 81 ± 1 | 74 ± 1 | 56 ± 1 | 89 ± 0 |
| | *higher depth (10+)* | 60 ± 1 | 56 ± 1 | 45 ± 1 | 80 ± 1 |
| *MAVE-NN_bb* | *shallow (5-9)* | 89 ± 4 | 74 ± 4 | 55 ± 2 | 88 ± 1 |
| | *higher depth (10+)* | 75 ± 11 | 57 ± 10 | 41 ± 13 | 82 ± 7 |
| *METL-global_1D* | *shallow (5-9)* | 86 ± 1 | 75 ± 1 | 55 ± 1 | 88 ± 1 |
| | *higher depth (10+)* | 63 ± 2 | 56 ± 1 | 42 ± 2 | 80 ± 2 |
| *METL-global_3D* | *shallow (5-9)* | 89 ± 1 | 77 ± 1 | 56 ± 2 | **90 ± 1** |
| | *higher depth (10+)* | 72 ± 3 | 59 ± 4 | **49 ± 4** | **89 ± 2** |
| *PT5-LoRA_reg* | *shallow (5-9)* | **93 ± 4** | **80 ± 4** | **60 ± 7** | **90 ± 4** |
| | *higher depth (10+)* | 81 ± 19 | 62 ± 10 | 46 ± 13 | 84 ± 17 |
| *PT5-LoRA_rank* | *shallow (5-9)* | **93 ± 2** | **80 ± 2** | 59 ± 3 | **90 ± 1** |
| | *higher depth (10+)* | **84 ± 7** | **65 ± 9** | 48 ± 8 | 87 ± 5 |
| *PT5-Emb* | *shallow (5-9)* | 92 ± 2 | 77 ± 4 | 53 ± 6 | 88 ± 3 |
| | *higher depth (10+)* | 77 ± 5 | 57 ± 9 | 46 ± 14 | 86 ± 6 |
| *PT5-Emb_evo* | *shallow (5-9)* | 92 ± 2 | 78 ± 1 | 56 ± 3 | **90 ± 1** |
| | *higher depth (10+)* | 76 ± 14 | 60 ± 11 | 47 ± 18 | 85 ± 11 |

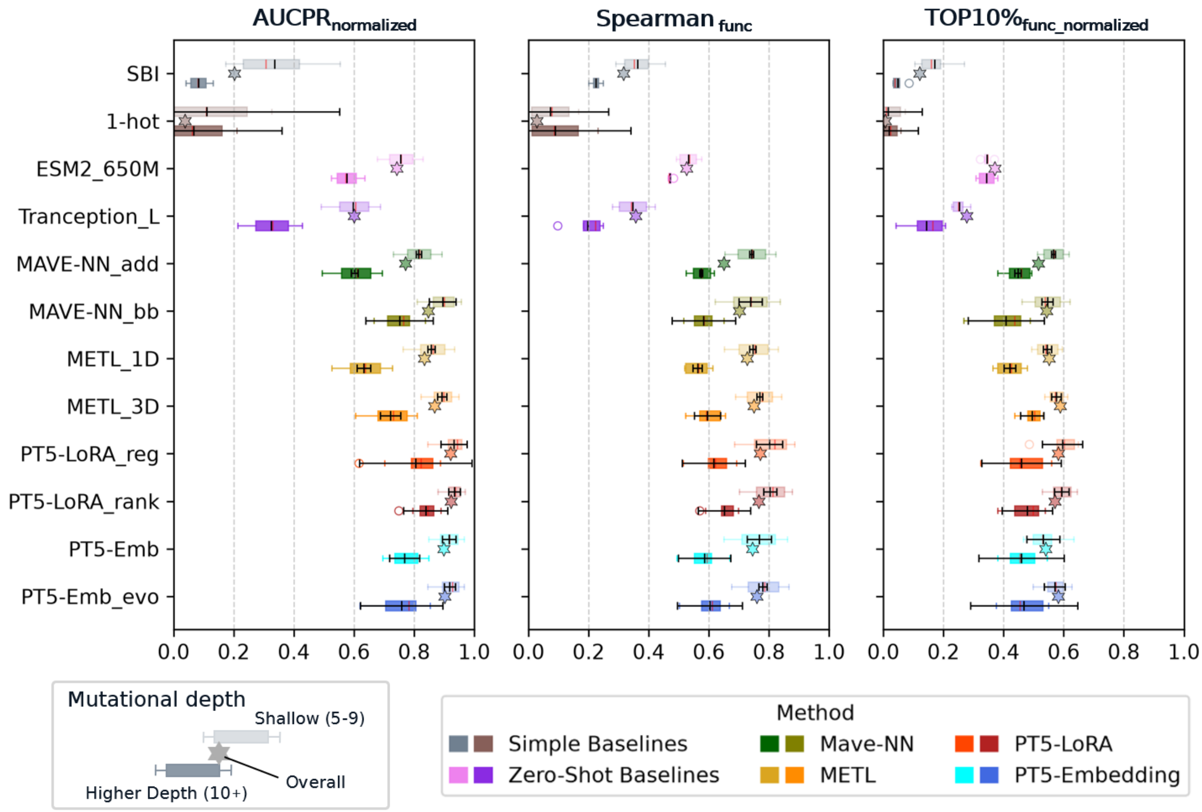


**Fig. S5: Phototropin results** †

### 6. SOM 6: Fine-tuning initialization

Instead of using the standard pretrained model with randomly initialized LoRA adapters, we investigated two alternative initialization approaches here.

For the Evo-tuned version we continued ProtT5's unsupervised pre-training on a set of homolog sequences to the protein of interest (we used the same model checkpoint we chose for embedding extraction for the "PT5-Emb evo" results in Table 1). Fine-tuning was started from this model checkpoint.

EVA[1] initializes LoRA weights based on the fine-tuning data to accelerate and increase performance of task specific fine-tuning.

When testing both in combination with our simple binary ranking loss for "LoRA_rank" (Table S11), we did not find any significant differences (Table S11). The "EVA initialized" version, was numerically better for the "shallow" depth while generalization at "higher depth" decreased, which was the opposite of desired behavior hinting at increased overfitting tendency.

---

**Table S11: DeePEn results for different fine-tuning initialization**[Δ]

| *Model* | *Depth* | $\mathit{AUCPR}_{norm}$ | $\mathit{Spearman}_{func}$ | $\mathit{TOP10\%}_{func}$ $_{norm}$ | $\mathit{NDCG@10\%}_{func}$ $_{norm}$ |
|---|---|---|---|---|---|
| *PT5-LoRA_rank* | *shallow (5-9)* | **79 ± 5** | 54 ± 4 | 34 ± 6 | **75 ± 10** |
| | *higher depth (10+)* | **64 ± 11** | **40 ± 10** | **27 ± 8** | 56 ± 44 |
| *PT5-LoRA_rank* | *shallow (5-9)* | **79 ± 5** | **55 ± 4** | **35 ± 4** | 74 ± 12 |
| *Evo-tuned* | *higher depth (10+)* | **64 ± 14** | 38 ± 5 | **27 ± 8** | **59 ± 37** |
| *PT5-LoRA_rank* | *shallow (5-9)* | **79 ± 2** | **55 ± 3** | **35 ± 5** | **75 ± 7** |
| *EVA initialized* | *higher depth (10+)* | 59 ± 8 | 39 ± 9 | 26 ± 7 | 47 ± 18 |

Δ Results per model for both depth bins (shallow/higher) give averages over all depth subsets and all four datasets. To control for shifting class prevalence, metrics are normalized (per depth subset) to a range between 0 (random) and 1 (perfect prediction). $\mathrm{Spearman}_{func}$ is not affected as its random performance is already 0. 95% confidence intervals of the mean values were only calculated for model replicates within individual depths/dataset-combinations. Metric values multiplied by 100, numerically best performing model per depth and metric in bold, these model differences are not statistically significant.

### 7. SOM 7: Comparing evo- and fine-tuned embeddings

By visualizing the embedding space (Fig. S6) we hoped for some additional insight on why using evo-tuned model embeddings of ProtT5 was better than using the pretrained model (Table 1, "PT5-Emb_evo" was better than "PT5-Emb"), while at the same time the use of the evo-tuned model as fine-tuning starting point had no positive effect (Table S11). One clear difference was the magnitude of change in the output embedding when comparing the base model output (blue) with evo-tuned embeddings (purple) and fine-tuned embeddings (red). Evo-tuning led to a slight shift while fine-tuning completely changed the embedding distribution. Also, per design fine-tuned model output showed clear fitness (DMS score) gradient while base model output and evo-tuned embeddings did not visibly show that.

We concluded from this that the starting point did not matter for fine-tuning because the changes introduced during the fine-tuning process are much larger than differences between starting points and initialization becomes less relevant. As others [2] have previously seen benefit from evo-tuning before fine-tuning we suspect that the high data regime present in DeePEn was the different maker. Even for the smallest training dataset, *Phototropin,* shown here (Fig. S6), we utilized over 3000 datapoints for training (Table 2).

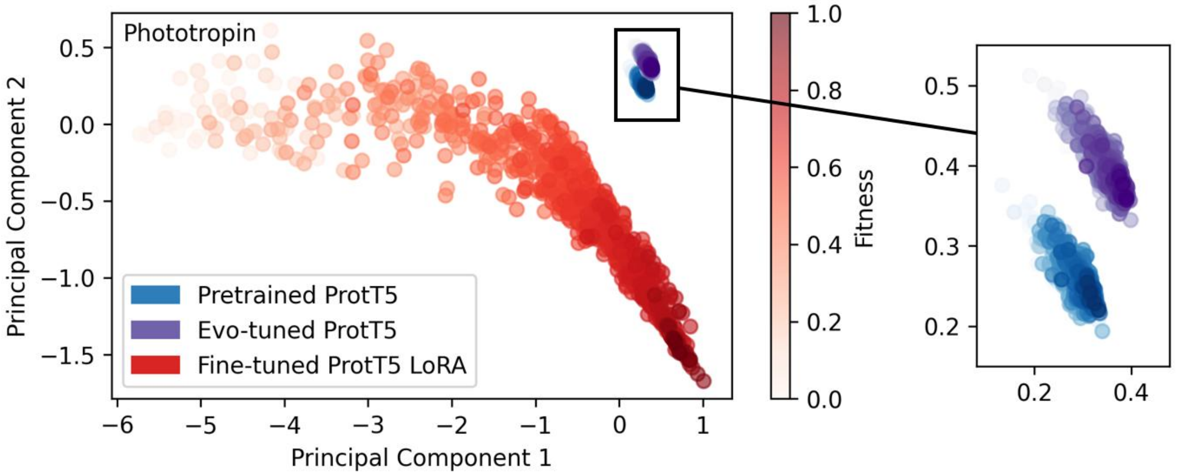


**Fig. S6: Embedding space changes.** PCA analysis of ProtT5 embeddings for a subset (1000 randomly drawn) of training sequences. The Fine-tuned (red) model showed a wide fitness aware embedding variance. Pretrained (blue) and Evo-tuned (purple) showed much less embedding variance and no visible fitness gradient. As both the Evo-tuned and Fine-tuned models originated from the Pretrained model state we concluded that $\Delta_{\text{Fine-tuning}} \gg \Delta_{\text{Evo-tuning}}$.

### 8. SOM 8: Ranking loss selection

To select the best performing loss for training, we tried out several versions of our ProtT5, LoRA fine-tuned prediction model. We left the prediction head unchanged and ran every model training three times with different random seeds. Metric calculation was done analog to results in Fig. 3 and Table 1. The *LoRA_reg* model used a standard MSE loss. In contrast we deployed a simple binary ranking loss for *LoRA_rank*. For each pair in a batch, we calculated the probability $P_{ij}$ of $label_i$ being greater than $label_j$ assuming gaussian noise with standard deviation sigma.

$$P_{ij} = P(label_i > label_j) = \Phi[(label_i - label_j) / (\sigma\sqrt{2})]$$

Where $\Phi$ is the cumulative distribution function (CDF) of the standard normal distribution. We used a StandardScaler to normalize the labels (scaled to mean of 0 and standard deviation of 1) and $\sigma = 0.05$ for the experimental noise. We then trained the model using binary cross entropy (BCE) loss on the model logit outputs for i and j trying to learn the probabilistic soft-labels $P_{ij}$:

$$loss_{BCE} = torch.nn.BCEWithLogitsLoss(output_i - output_j, P_{ij})$$

Adding on top of the *LoRA_rank* version we further complemented the loss with a contrastive loss calculated based on the embeddings of i and j.

$$loss_{contrastive} = (1 - target) * \left(1 - similarity_{ij}\right)^2 + target * ReLU\ (similarity_{ij} - margin)^2$$

with

$$target = (\left(P_{ij} - 0.5\right) * 2)^2$$

With $similarity_{ij}$ being the cosine similarity between their embeddings and target being 0 for similar $label_i$ and $label_j$ and 1 for dissimilar ones based on $P_{ij}$. Both losses were added up and we weighed the contrastive loss 10 times stronger than the BCE loss. The resulting model was dubbed LoRA_rank_contrastive. For the second contrastive version *LoRA_rank_contrastive_onesided* we left out the first half of the sum for the contrastive loss to only punish very similar embeddings with dissimilar fitness values but allowed the reverse.

With the last variant *LoRA_rank_lambda_rank* we optimized the NDCG following previous work [3]. We set the necessary relevance values in the same way we defined them for the evaluation metric $NDCG@10\%_{func}$. By setting the relevance score to 0 for all non-functional variants and then using linear min-max scaling of the continuous DMS scores to assign values between 1 (lowest functional score) and 3 (highest functional score) to functional variants. We also only calculated the loss on pairs of values with a $delta_{ij}$ of at least 0.2 (on normalized values) to avoid potential training on any mislabeled pairs.

While the differences between different losses were insignificant, Table S12 showed clearly that the ranking models came out on top numerically compared to the regression version. We selected the lambda_rank to be the *LoRA_rank* version for the main comparison as it performed best out of the different ranking variants.

#### Table S12: DeePEn results for different ranking losses[Δ]

| *Model* | *Depth* | $AUCPR_{norm}$ | $Spearman_{func}$ | $TOP10\%_{func,norm}$ | $NDCG@10\%_{func,norm}$ |
|---|---|---|---|---|---|
| *PT5-LoRA_reg* | *shallow (5-9)* | 81 ± 5 | 53 ± 4 | 33 ± 6 | 76 ± 11 |
| | *higher depth (10+)* | 65 ± 16 | 36 ± 8 | 25 ± 13 | 57 ± 27 |
| *PT5-LoRA_rank* | *shallow (5-9)* | 79 ± 5 | **54 ± 4** | **34 ± 6** | 75 ± 10 |
| | *higher depth (10+)* | 64 ± 11 | **40 ± 10** | 27 ± 8 | 56 ± 44 |
| *PT5-LoRA_rank contrastive* | *shallow (5-9)* | 79 ± 5 | **54 ± 5** | **34 ± 7** | 75 ± 11 |
| | *higher depth (10+)* | 62 ± 9 | **40 ± 7** | **28 ± 8** | 55 ± 23 |
| *PT5-LoRA_rank contrastive_onesided* | *shallow (5-9)* | 79 ± 2 | 53 ± 4 | 33 ± 6 | 74 ± 4 |
| | *higher depth (10+)* | 62 ± 10 | 39 ± 14 | 24 ± 11 | 50 ± 26 |
| *PT5-LoRA_rank lambda_rank* | *shallow (5-9)* | **82 ± 3** | **54 ± 4** | 33 ± 7 | **82 ± 4** |
| | *higher depth (10+)* | **68 ± 15** | 39 ± 13 | 27 ± 10 | **69 ± 25** |

[Δ] Results per model for both depth bins (shallow/higher) give averages over all depth subsets and all four datasets.To control for shifting class prevalence, metrics are normalized (per depth subset) to a range between 0 (random) and 1 (perfect prediction). $Spearman_{func}$ is not affected as its random performance is already 0. 95% confidence intervals of the mean values were only calculated for model replicates within individual depths/dataset-combinations. Metric values multiplied by 100, numerically best performing model per depth and metric in bold, these model differences are not statistically significant.

### 9. SOM 9: DeePEn results for TOPn%$_{func}$-Recall

We provided the TOPn%$_{func}$ for 6 different thresholds in Table S13. Both *PT5-LoRA* models were numerically on top across thresholds. Naturally it was harder to achieve high performance for lower thresholds, which was also reflected in the *Random* and normalized *Perfect classifier* baseline. The ranking was mostly consistent across the range of thresholds, which makes the threshold selection less crucial. We went with TOP10%$_{func}$ for the main manuscript to match the NDCG@10% threshold and focus specifically on the very top of variants.

---

**Table S13: TOPn results**[Δ]

| *Model* | *Depth* | *TOP5%$_{func}$* | *TOP10%$_{func}$* | *TOP15%$_{func}$* | *TOP20%$_{func}$* | *TOP25%$_{func}$* | *TOP30%$_{func}$* |
|---|---|---|---|---|---|---|---|
| *Random baseline* | *shallow (5-9)* | 2 ± 0 | 5 ± 0 | 7 ± 0 | 10 ± 0 | 12 ± 0 | 15 ± 0 |
| | *hi. depth (10+)* | 1 ± 0 | 3 ± 0 | 5 ± 0 | 6 ± 0 | 8 ± 0 | 9 ± 0 |
| *Model* | *Depth* | *TOP5%$_{func}$ norm* | *TOP10%$_{func}$ norm* | *TOP15%$_{func}$ norm* | *TOP20%$_{func}$ norm* | *TOP25%$_{func}$ norm* | *TOP30%$_{func}$ norm* |
| *Perfect classifier baseline* | *shallow (5-9)* | 3 ± 0 | 5 ± 0 | 8 ± 0 | 11 ± 0 | 14 ± 0 | 17 ± 0 |
| | *hi. depth (10+)* | 3 ± 0 | 7 ± 0 | 11 ± 0 | 15 ± 0 | 18 ± 0 | 22 ± 0 |
| *SBI baseline* | *shallow (5-9)* | 7 ± 0 | 10 ± 0 | 13 ± 0 | 14 ± 0 | 15 ± 0 | 16 ± 0 |
| | *hi. depth (10+)* | 3 ± 0 | 3 ± 0 | 3 ± 0 | 3 ± 0 | 4 ± 0 | 4 ± 0 |
| *1-hot baseline* | *shallow (5-9)* | 0 ± 3 | 1 ± 5 | 1 ± 6 | 2 ± 7 | 2 ± 9 | 3 ± 9 |
| | *hi. depth (10+)* | 0 ± 4 | 0 ± 5 | 1 ± 6 | 2 ± 9 | 2 ± 10 | 3 ± 12 |
| *ESM2_650M$_{zero\text{-}shot}$* | *shallow (5-9)* | 9 ± 0 | 13 ± 0 | 16 ± 0 | 17 ± 0 | 19 ± 0 | 20 ± 0 |
| | *hi. depth (10+)* | 12 ± 0 | 17 ± 0 | 20 ± 0 | 22 ± 0 | 25 ± 0 | 28 ± 0 |
| *Tranception_L$_{zero\text{-}shot}$* | *shallow (5-9)* | 10 ± 0 | 13 ± 0 | 16 ± 0 | 19 ± 0 | 22 ± 0 | 26 ± 0 |
| | *hi. depth (10+)* | 7 ± 0 | 15 ± 0 | 18 ± 0 | 20 ± 0 | 23 ± 0 | 25 ± 0 |
| *MAVE-NN_add* | *shallow (5-9)* | 21 ± 2 | 25 ± 2 | 28 ± 2 | 33 ± 1 | 35 ± 1 | 38 ± 2 |
| | *hi. depth (10+)* | 19 ± 2 | 22 ± 2 | 20 ± 0 | 20 ± 2 | 21 ± 1 | 22 ± 2 |
| *MAVE-NN_bb* | *shallow (5-9)* | 23 ± 11 | 27 ± 12 | 30 ± 12 | 33 ± 12 | 36 ± 13 | 39 ± 12 |
| | *hi. depth (10+)* | 13 ± 5 | 20 ± 8 | 20 ± 7 | 22 ± 6 | 25 ± 8 | 25 ± 11 |
| *METL-global_1D* | *shallow (5-9)* | 24 ± 5 | 28 ± 4 | 32 ± 3 | 35 ± 4 | 38 ± 3 | 40 ± 3 |
| | *hi. depth (10+)* | 13 ± 4 | 19 ± 6 | 21 ± 6 | 23 ± 5 | 26 ± 6 | 28 ± 5 |
| *METL-global_3D* | *shallow (5-9)* | 25 ± 6 | 28 ± 3 | 32 ± 4 | 35 ± 4 | 37 ± 3 | 39 ± 4 |
| | *hi. depth (10+)* | 16 ± 5 | 25 ± 6 | 25 ± 5 | 27 ± 7 | 30 ± 5 | 32 ± 6 |
| *PT5-LoRA_reg* | *shallow (5-9)* | **28 ± 8** | **33 ± 6** | **37 ± 6** | **40 ± 4** | 42 ± 5 | 45 ± 5 |
| | *hi. depth (10+)* | 19 ± 6 | 25 ± 13 | **28 ± 12** | **32 ± 15** | 34 ± 12 | 35 ± 14 |
| *PT5-LoRA_rank* | *shallow (5-9)* | **28 ± 4** | **33 ± 7** | 36 ± 7 | **40 ± 5** | **43 ± 4** | **46 ± 4** |
| | *hi. depth (10+)* | **24 ± 8** | **27 ± 10** | **28 ± 14** | **32 ± 11** | **35 ± 13** | **37 ± 14** |
| *PT5-Emb* | *shallow (5-9)* | 21 ± 7 | 25 ± 5 | 28 ± 6 | 33 ± 6 | 37 ± 5 | 40 ± 5 |
| | *hi. depth (10+)* | 14 ± 12 | 19 ± 12 | 21 ± 9 | 25 ± 9 | 27 ± 7 | 30 ± 8 |
| *PT5-Emb_evo* | *shallow (5-9)* | 25 ± 10 | 28 ± 7 | 32 ± 6 | 36 ± 4 | 40 ± 4 | 44 ± 4 |
| | *hi. depth (10+)* | 17 ± 9 | 21 ± 16 | 24 ± 14 | 28 ± 16 | 31 ± 17 | 34 ± 18 |

[Δ] Results per model for both depth bins show each respective metric averaged across depth subsets. Absolute random baseline values in the top row, followed by baseline methods, and supervised advanced methods below. To control for shifting class prevalence, metrics are normalized (per depth subset) to a range between 0 (random) and 1 (perfect prediction). 95% confidence intervals of the mean values were only calculated for model replicates within individual depths/dataset-combinations. Metric values multiplied by 100, numerically best performing model per depth and TOPn threshold in bold, these model differences are not statistically significant. The *Random baseline* is randomly ranking all variants, while the *Perfect classifier baseline* is perfectly separating non-functional (at the bottom) and functional variants (at the top) while randomly shuffling all functional variants.

### 10. SOM 10: DeePEn model ranking consistency across metrics

We analyzed correlations between the different evaluation metrics across different models, again we used the normalized metrics. For this we aggregated results slightly differently than for the benchmark itself. Here we did not average across training runs (random seeds) but binned results of individual training runs directly into the known shallow and higher depth distance bins (Table S3-S6), as this allowed us to have more statistical power (30 training run averages instead of 12 model averages) for this analysis. We then calculated correlations between model (training run) performances for different metrics. We found moderate to high Spearman correlations (Fig. S7). Especially for the shallow subset (blue/green), this means that all four metrics captured some unique aspects of model performance and therefore provided additional information. Interestingly in both depth bins AUCPR and NDCG10%$_{func}$ were highly correlated which points towards the fact that NDCG is highly influenced by binary classification quality (false positives are very punishing because non-functional variants have a relevance score of 0, while functional variants have a score of 1-3).

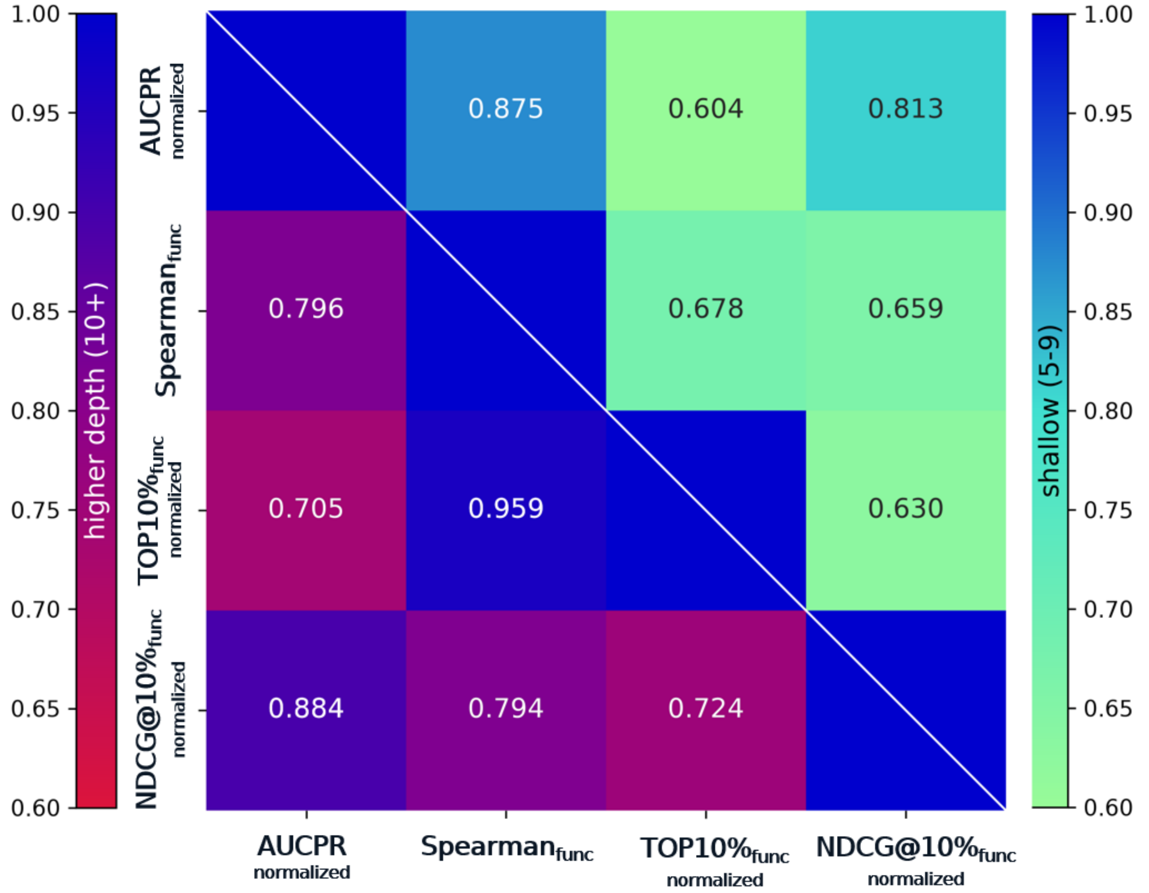


**Fig. S7: DeePEn metric correlations.** Spearman correlation between the four selected evaluation metrics. Top right (blue green): on the shallow depth bin Bottom left (Blue red): on the higher depth bin. Correlations were calculated on individual training run results instead of averaged (across random seeds) model results. We found moderately high correlations, which hinted at unique information content for all metrics.

Next, we looked at model ranking. Fig. S8 shows this for AUCPR, Spearman$_{func}$ and TOP10%$_{func}$. The vertical method order is the average rank per method. As seen in the main Fig. 3 / Table 1 results the LoRA_rank model is on the top. But in line with the dataset specific results (Table S7-S10 and Fig. S2-S5) there are some outliers.
Both zero-shot baselines (Tranception_L and ESM2_650M) overperformed on the *Dehydratase* dataset, especially at higher depth. This makes sense as the dataset is recombining different cross-species dehydratase wildtypes (on which evolutionary zero-shot models are well informed on), resulting in a dataset where many higher order variants (nearly 50%, see SOM 11) are not combinations of lower order known variants. This apparently made it harder to predict, for the supervised methods.
The second outlier are both MAVE-NN models on the GFP dataset. We suspect that the weak epistatic effects (Fig. S9) of this dataset align well with the underlying additive assumption of their model design.
One important limitation is that model differences were often small and statistically not significant, which makes conversion of those differences into absolute ranks less reliable. Global trends hold (top methods are better than bottom ones) but local order between two neighboring methods will not be statistically robust.

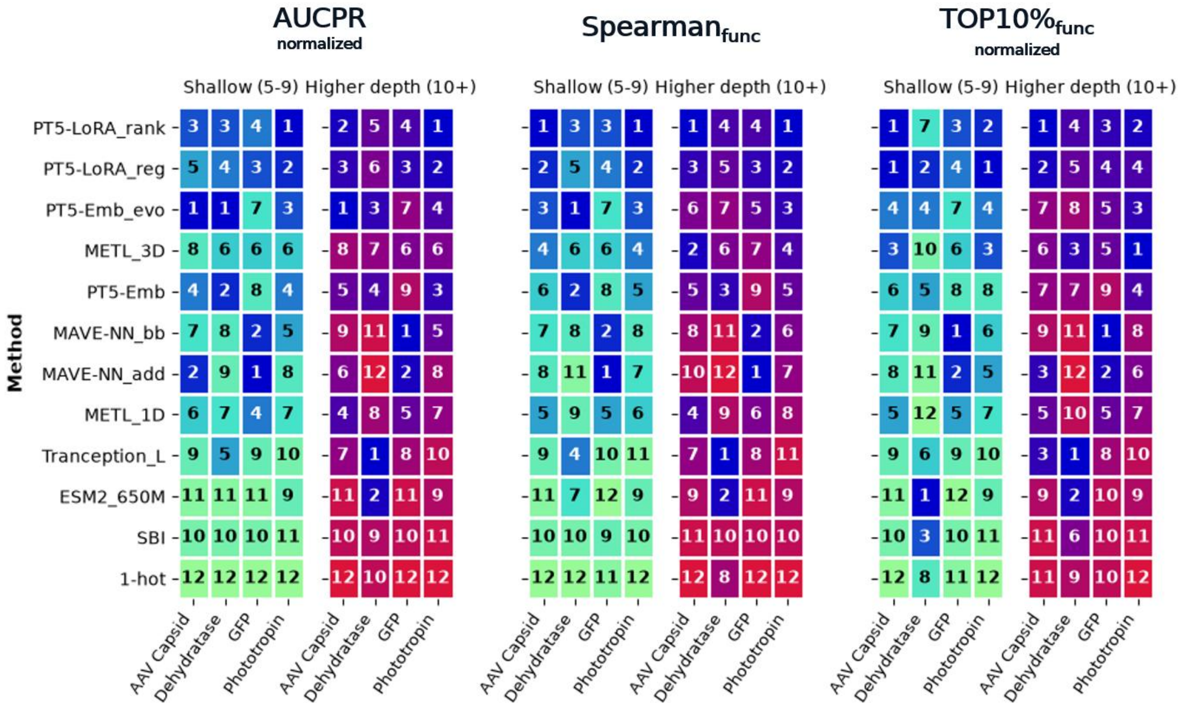


**Fig. S8: DeePEn method ranking.** Method ranking per metric (subplots) and dataset (x-axis). Left (Blue green): on the shallow depth bin, Right (Blue red): on the higher depth bin. Method rankings were calculated on normalized subset results, average per bin for individual datasets. Method order on the y axis is determined by the sum of the entire row (lower sum/better method is higher up).

### 11. SOM 11: Epistasis

We analysed the epistatic interactions for all four DeePEn datasets (Fig. S9). We followed the method used by Chen et al. in their analysis of the *Phototropin* data[4]. The authors utilized an additive model on logarithmic scale for epistasis, where $F_{comb}$, $F_{single}$ and $F_{wt}$ are the log fitness values (DMS scores) for higher order recombinants, single mutants and the wildtype sequence respectively. *ProteinGym* DMS_scores for the four datasets were logarithmic already with exception of the *Phototropin* dataset. Therefore, we went back to the original source data[4] and collected the missing log values for this dataset.

$$epistasis = (F_{comb} - F_{wt}) - \sum(F_{single} - F_{wt})$$

Positive epistasis values signal synergistic effect accumulation of single mutant effects, while negative values indicate an anti-synergistic interaction. We defined weak epistasis for all cases where the deviation from expected additive effect was within experimental uncertainty (calculated based on the three available replicates of the *Phototropin* dataset and scaled for the other datasets).

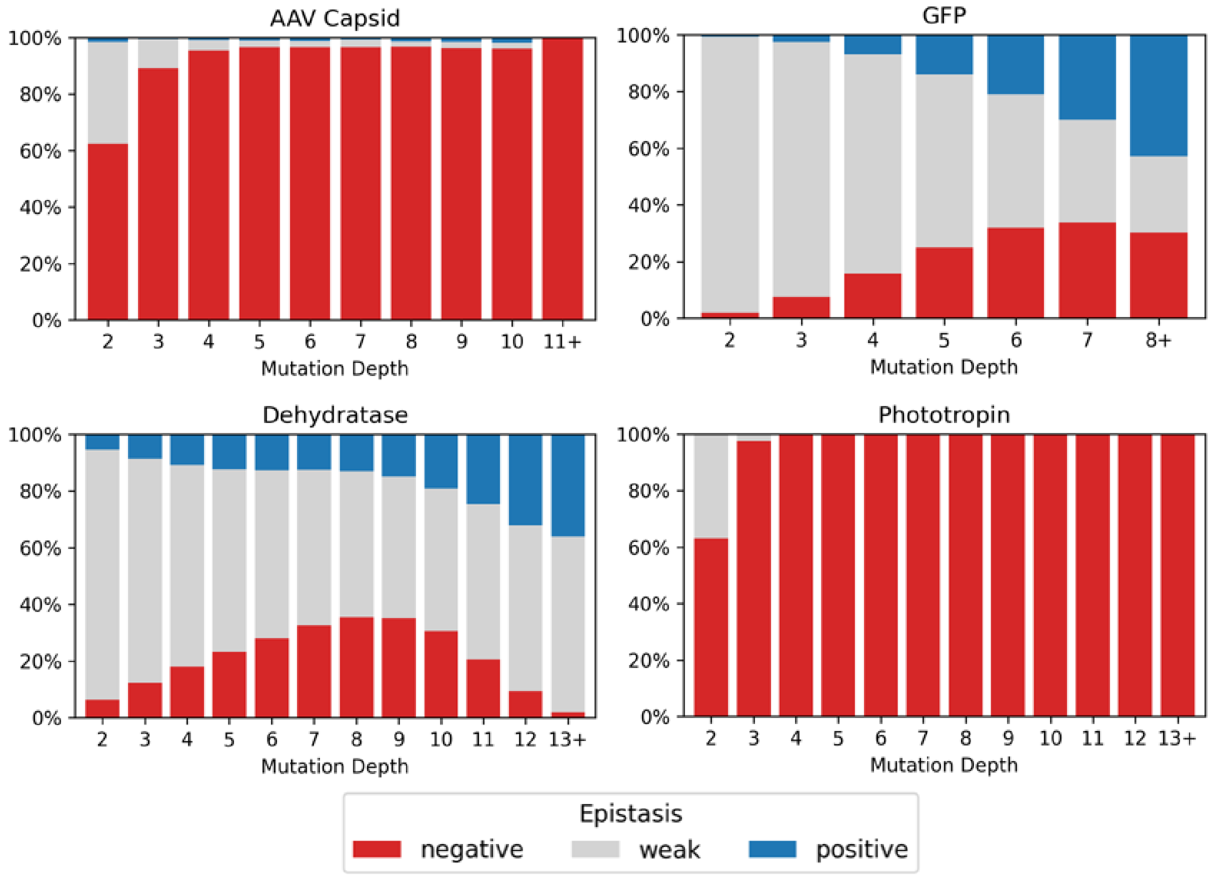


**Fig. S9: Epistasis analysis.** For each of the four DeePEn datasets we provide an epistatic analysis by mutation depth (x-axis). Negative epistasis (red) signals conflicting interaction (less than additive) while positive (blue) signals synergies. For weak epistasis (grey) the effect of the recombinant variant was close to the expected (additive) effect across all involved single substitutions. We used all fitness values on logarithmic scale for this analysis.

We found that the randomly sampled datasets *GFP* and *Dehydratase* showed a lot of weak and even some synergistic epistatic interaction while the other two dataset *AAV capsid* (which originates from an in-silico engineering campaign) and *Phototropin* (which was created using rational design, i.e. combining functional singles to create highly active recombinants) were dominated by negative epistasis. In general, this hints at limitations of rational design, when it comes to more distant recombinant designs.
We had to exclude higher order variants if not all single amino acid substitutions were available. This occurred in the *GFP* dataset, where we could not analyze 7.959 variants (15.7%) and more severely in the *Dehydratase* set where we couldn't evaluate nearly half (48.7%, 241.355 variants).

### 12. SOM 12: Sampling functional variants

Here we investigated the usefulness of model predictions for pre-selection of variants to experimentally validate. We compared our PT5-LoRA_rank model with the blackbox *MAVE-NN_bb* version and *METL_3D*. For each test set variant we calculated an average predicted rank (from the three training runs with different seeds) per model. We then selected variants from the top of this ranking until we reached 95% recall of all functional variants (Fig. S10a) / 95% recall of the top 10% of functional variants (Fig. S10b). We reported the precision and the percentage of the dataset we needed to sample for each model and dataset. If only functional variants are of interest (Fig. S10a), between 29% (for *Dehydratase*) and 79% (for *GFP*) of the experimental effort can be saved while still identifying 95% of functional variants. Precision was high for all datasets, but the maximum potential saving depends on the share of functional variants in each set. Likewise, if only the most active variants are desired (Fig. S10b), even less sampling would be needed. Precision was a lot lower here, but for the *AAV Capsid* and *Phototropin* datasets some sampling efficiency could be gained. In contrast the models' bad ranking of functional *Dehydratase* variants (compare Spearman$_{func}$ in Fig. S3 and Table S8) would not allow to sample much less.

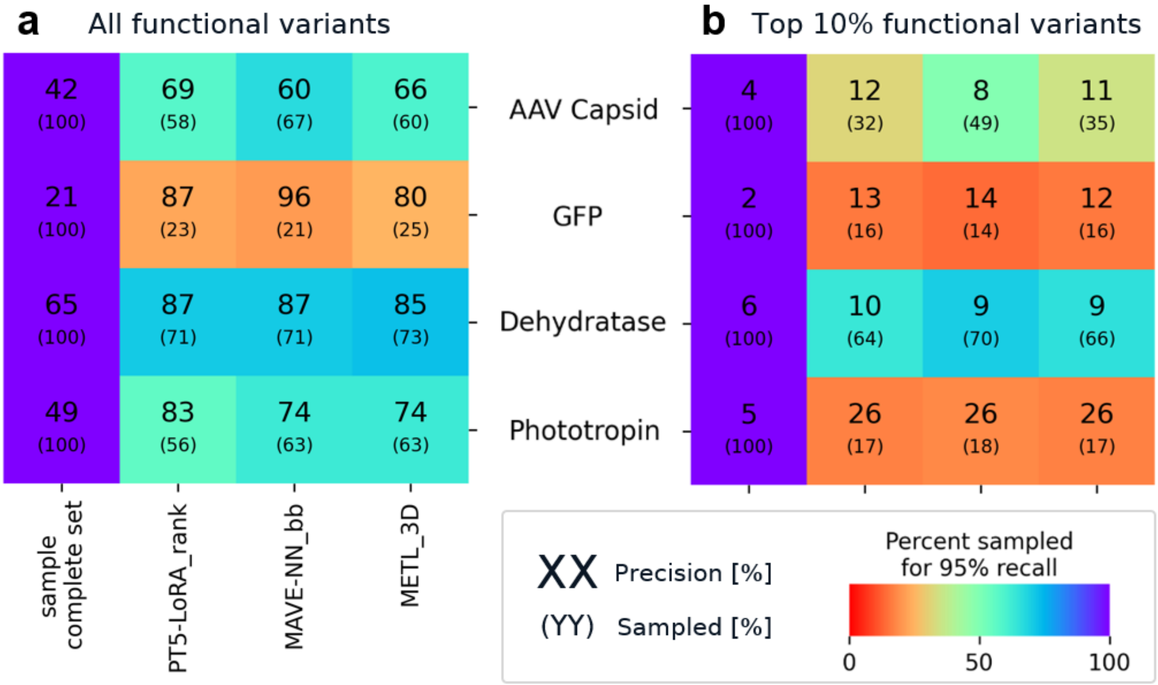


**Fig. S10: Sampling efficiency for 95% recall.** We computed the average predicted rank (from three training runs with different random seeds) for all models (x-axis). We then sampled from the top until we reached 95% recall for all functional variants (Panel **a**) / the top 10% functional variants (Panel **b**). We reported the precision and the share of the overall dataset that needs to be sampled (coloring is also showing this share). In the left column we provided complete sampling (Sampled 100%) of the set as a baseline, to provide information about the prevalence of all / top10% functional variants (Precision).

One inherent limitation is that the variant effects of depth 1-3 (and at least some amount of depth 4 variants as a validation set) need to be known to realize those efficiency gains, which splits the experimental campaign into two separate steps. The potential gains that can be realized by this approach will also be heavily influenced by the prevalence of functional sequences in your dataset.

Proteins with narrow fitness landscapes like the *GFP* example shown here, with only 21% functional variants, naturally show higher potential gains. Another important impact factor is the selection method for investigated sequences, random mutagenesis (*GFP*), random recombination of orthologous sequences (*Dehydratase*), selective recombination based on full single amino acid mutagenesis (*Phototropin*) and ML guided selection (*AAV Capsid*) produce vastly different datasets with different characteristics and challenges.

### 13. SOM 13: Binarization threshold sensitivity

Instead of using the ProteinGym binarization threshold, here we investigated the model performance when the wildtype is used as threshold. As producing variants better than wildtype is often a key part of engineering efforts we wanted to see how sensitive the model performance is to this change. Below (Tables S14-S17, caption at Table S14) we are comparing the subset normalized AUCPR of the PT5-LoRA_reg model (we did not use the better PT5-LoRA_rank model here as the binarization was used for calculating relevance scores during it's training) for the wildtype binarized and the ProteinGym binarized setting. The *AAV Capsid* dataset stood out showing little difference. This was due to the ProteinGym threshold being close to the wildtype already. For the other three datasets the model's discriminative power was much lower when using it as a "better than wildtype"-classifier. This supports our view that using those models to filter out non-functional sequences enables efficiency gains (Fig. S10), but identifying highly functional subpopulations remains much harder (main Fig. 6 and discussion).

---

**Table S14: AAV capsid - wildtype binarization** ^Δ^

| ProteinGym binarization | | | | Wildtype binarization | | | | |
|---|---|---|---|---|---|---|---|---|
| Distance from wildtype | Number of seq | Number of seq >thresh. | AUCPR norm | Distance from wildtype | Number of seq | Number of seq >thresh. | AUCPR norm | ΔAUCPR norm |
| 5 | 6,298 | 3,772 | 84.2% | 5 | 6,298 | 3,673 | 82.6% | -1.6% |
| 6 | 5,396 | 2,584 | 71.9% | 6 | 5,396 | 2,481 | 70.1% | -1.8% |
| 7 | 858 | 398 | 69.7% | 7 | 858 | 379 | 66.4% | -3.3% |
| 8 | 739 | 237 | 53.2% | 8 | 739 | 221 | 45.3% | -7.9% |
| 9 | 651 | 141 | 39.5% | 9 | 651 | 130 | 37.3% | -2.2% |
| 10 | 583 | 92 | 35.5% | 10 | 583 | 85 | 31.5% | -4.0% |
| 11-28 | 2,886 | 91 | 15.8% | 11-28 | 2,886 | 77 | 14.5% | -1.3% |

Δ Normalized AUCPR values for the PT5-LoRA_reg model. We report average depth wise results to make the shift in class prevalence visible. Due to this prevalence shift, we extended the last subset to include shallower depth when it would otherwise contain less than 50 variants above the threshold. Notable changes in red.

---

**Table S15: GFP - wildtype binarization** ^Δ^

| ProteinGym binarization | | | | Wildtype binarization | | | | |
|---|---|---|---|---|---|---|---|---|
| Distance from wildtype | Number of seq | Number of seq >thresh. | AUCPR norm | Distance from wildtype | Number of seq | Number of seq >thresh. | AUCPR norm | ΔAUCPR norm |
| 5 | 6,825 | 2,284 | 97.0% | 5 | 6,825 | 238 | 29.9% | -67.1% |
| 6 | 4,298 | 802 | 94.7% | | | | | -64.2% |
| 7 | 2,526 | 229 | 91.0% | 6-15 | 9,305 | 96 | 30.5% | -60.5% |
| 8-15 | 2,481 | 73 | 80.9% | | | | | -50.4% |

#### Table S16: Dehydratase - wildtype binarization [Δ]

| ProteinGym binarization | | | | Wildtype binarization | | | | |
|---|---|---|---|---|---|---|---|---|
| Distance from wildtype | Number of seq | Number of seq >thresh. | AUCPR norm | Distance from wildtype | Number of seq | Number of seq >thresh. | AUCPR norm | ΔAUCPR norm |
| 5 | 60,263 | 45,987 | 89.9% | 5 | 60,263 | 18,219 | 25.4% | -64.5% |
| 6 | 98,869 | 70,269 | 85.3% | 6 | 98,869 | 25,967 | 22.8% | -62.5% |
| 7 | 115,751 | 76,096 | 78.2% | 7 | 115,751 | 25,064 | 18.9% | -59.3% |
| 8 | 97,639 | 58,953 | 70.5% | 8 | 97,639 | 17,664 | 15.7% | -54.8% |
| 9 | 57,139 | 32,216 | 63.2% | 9 | 57,139 | 8,753 | 12.7% | -50.5% |
| 10 | 22,942 | 12,488 | 54.4% | 10 | 22,942 | 3,374 | 10.2% | -44.2% |
| 11 | 6,370 | 3,727 | 55.2% | 11 | 6,370 | 1,024 | 9.2% | -46.0% |
| 12 | 1,354 | 920 | 69.7% | 12-28 | 1,967 | 303 | 13.8% | -55.9% |
| 13-28 | 613 | 173 | 79.0% | | | | | -65.2% |

#### Table S17: Phototropin - wildtype binarization [Δ]

| ProteinGym binarization | | | | Wildtype binarization | | | | |
|---|---|---|---|---|---|---|---|---|
| Distance from wildtype | Number of seq | Number of seq >thresh. | AUCPR norm | Distance from wildtype | Number of seq | Number of seq >thresh. | AUCPR norm | ΔAUCPR norm |
| 5 | 9,603 | 7,015 | 97.0% | 5 | 9,603 | 2,403 | 87.5% | -9.5% |
| 6 | 19,193 | 12,518 | 95.3% | 6 | 19,193 | 3,688 | 83.2% | -12.1% |
| 7 | 29,255 | 16,938 | 93.6% | 7 | 29,255 | 4,279 | 76.5% | -17.1% |
| 8 | 34,230 | 17,071 | 91.4% | 8 | 34,230 | 3,669 | 70.9% | -20.5% |
| 9 | 30,925 | 13,010 | 88.5% | 9 | 30,925 | 2,453 | 65.3% | -23.2% |
| 10 | 21,167 | 7,361 | 85.8% | 10 | 21,167 | 1,188 | 59.6% | -26.2% |
| 11 | 11,022 | 3,057 | 82.5% | 11 | 11,022 | 432 | 56.3% | -26.2% |
| 12 | 4,093 | 905 | 78.6% | 12-15 | 5,293 | 109 | 45.5% | -33.1% |
| 13-15 | 1,200 | 182 | 75.1% | | | | | -29.6% |
